# Centrioles generate a local pulse of Polo/PLK1 activity to initiate mitotic centrosome assembly

**DOI:** 10.1101/2021.10.26.465695

**Authors:** Siu-Shing Wong, Zachary M. Wilmott, Saroj Saurya, Ines Alvarez-Rodrigo, Felix Y. Zhou, Kwai-Yin Chau, Alain Goriely, Jordan W. Raff

**Affiliations:** Sir William Dunn School of Pathology, University of Oxford, Oxford, UK; Mathematical Institute, University of Oxford, Oxford, UK; The Francis Crick Institute, London, UK; Ludwig Institute for Cancer Research, Nuffield Department of Clinical Medicine, University of Oxford, Oxford, UK. Current Address: University of Texas, Southwestern Medical Centre; Department of Computer Science, University of Bath, Bath, UK

**Author notes:** These authors contributed equally to this work (Siu-Shing Wong, Zachary M. Wilmott).

**Keywords:** centriole, centrosome, PCM, PLK1, mitosis, cell cycle, oscillator

## Abstract

Mitotic centrosomes are formed when centrioles start to recruit large amounts of pericentriolar material (PCM) around themselves in preparation for mitosis. This centrosome “maturation” requires the centrioles and also Polo/PLK1 protein kinase. The PCM comprises several hundred proteins and, in *Drosophila*, Polo cooperates with the conserved centrosome proteins Spd-2/CEP192 and Cnn/CDK5RAP2 to assemble a PCM scaffold around the mother centriole that then recruits other PCM client proteins. We show here that in *Drosophila* syncytial blastoderm embryos, centrosomal Polo levels rise and fall during the assembly process—peaking, and then starting to decline, even as levels of the PCM scaffold continue to rise and plateau. Experiments and mathematical modelling indicate that a centriolar pulse of Polo activity, potentially generated by the interaction between Polo and its centriole receptor Ana1 (CEP295 in humans), could explain these unexpected scaffold assembly dynamics. We propose that centrioles generate a local pulse of Polo activity prior to mitotic entry to initiate centrosome maturation, explaining why centrioles and Polo/PLK1 are normally essential for this process.

## Introduction

Centrosomes are important organisers of the cell that are formed when mother centrioles recruit a matrix of pericentriolar material (PCM) around themselves (Conduit *et al*, 2015; Bornens, 2021; Vasquez-Limeta & Loncarek, 2021; Lee *et al*, 2021; Woodruff, 2021). The PCM contains several hundred proteins (Alves-Cruzeiro *et al*, 2013), including many that help nucleate and organise microtubules (MTs), as well as many signalling molecules, cell cycle regulators, and checkpoint proteins. In this way, the centrosomes function as major MT organising centres (MTOC) and also important coordination centres in many cell types (Arquint *et al*, 2014; Chavali *et al*, 2014).

In interphase, most cells organise relatively little PCM, but there is a dramatic increase in PCM recruitment as cells prepare to enter mitosis—a process termed centrosome maturation (Palazzo *et al*, 2000; Conduit *et al*, 2015). Centrioles are required to initiate efficient mitotic PCM assembly (Bobinnec *et al*, 1998; Kirkham *et al*, 2003; Basto *et al*, 2006; Sir *et al*, 2013; Bazzi & Anderson, 2014; Wong *et al*, 2015), and, in worm embryos, centrioles are continuously required to promote the growth of the mitotic PCM—although they are not required to maintain the mitotic PCM once it has reached its full size (Cabral *et al*, 2019).

The protein kinase Polo/PLK1 is also required for the assembly of the mitotic PCM in many, if not all, systems (Sunkel & Glover, 1988; Lane & Nigg, 1996; Dobbelaere *et al*, 2008; Haren *et al*, 2009; Lee & Rhee, 2011; Conduit *et al*, 2014a; Woodruff *et al*, 2015b; Ohta *et al*, 2021). PLK1 performs many functions during mitosis (Archambault & Glover, 2009; Colicino & Hehnly, 2018), and it is recruited to different locations within the cell via its Polo-Box-Domain (PBD), which binds to phosphorylated S-S(P)/T(P) motifs on various scaffolding proteins (Elia *et al*, 2003; Reynolds & Ohkura, 2003; Song *et al*, 2000; Seong *et al*, 2002). Importantly, PBD binding to these scaffolding proteins helps to activate PLK1 by relieving an inhibitory interaction between the PBD and the kinase domain (Xu *et al*, 2013), although full activation also requires phosphorylation (Archambault & Glover, 2009; Colicino & Hehnly, 2018). PLK1 is recruited to centrosomes by the scaffolding protein CEP192 in vertebrates (Joukov *et al*, 2010, 2014; Meng *et al*, 2015), and by the CEP192 homologues Spd-2/SPD-2 in flies and worms (Decker *et al*, 2011; Alvarez-Rodrigo *et al*, 2019; Ohta *et al*, 2021). In these species, the Polo/PLK-1 recruited by Spd-2/SPD-2 can then phosphorylate Cnn/SPD-5 (flies/worms), which allows these large helical proteins to assemble into macromolecular PCM-scaffolds that help recruit the many other PCM “client” proteins (Conduit *et al*, 2014a; Woodruff *et al*, 2015a; Feng *et al*, 2017; Woodruff *et al*, 2017; Cabral *et al*, 2019; Ohta *et al*, 2021).

Here, we focus on the kinetics of mitotic PCM scaffold assembly in living *Drosophila* syncytial blastoderm embryos—where we can simultaneously track the behaviour of tens to hundreds of centrosomes as they rapidly and near-synchronously assemble over several nuclear division cycles that occur in a common cytoplasm. Surprisingly, we observe that the centrosomal levels of Polo rise and fall during the assembly process, with centrosomal levels peaking, and then starting to decline, even as the Cnn scaffold continues to grow. Mathematical modelling and further experiments indicate that an interaction between Polo and its centriole receptor Ana1 (CEP295 in vertebrates) could generate a local pulse of centriolar Polo activity, and that such a mechanism could explain the unexpected assembly kinetics of the PCM scaffold. We propose that centrioles generate a local pulse of Polo activity that initiates mitotic centrosome assembly in syncytial fly embryos prior to mitotic entry. We speculate that the ability of centrioles to locally activate Polo/PLK1 prior to mitosis may be a conserved feature of mitotic centrosome assembly—explaining why centrioles and Polo/PLK1 are both normally required to initiate this process.

## Results

### PCM-scaffold proteins exhibit distinct assembly dynamics

To better understand how Spd-2, Polo and Cnn cooperate to assemble the PCM scaffold we quantified their recruitment dynamics in syncytial *Drosophila* embryos during nuclear cycles 11-13 (Figure 1). Note that we have not attempted to quantify (nor model—see below) the dramatic disassembly of the mitotic PCM that occurs at the end of mitosis, as in fly and worm embryos this is a complicated process in which large “packets” or “flares” of the mitotic PCM are mechanically removed from the PCM in a MT-dependent manner (Megraw *et al*, 2002; Magescas *et al*, 2019; Mittasch *et al*, 2020). In the experiments reported here we used fluorescent reporters fused to several different fluorescent tags—Neon Green (NG), GFP, RFP or mCherry—and expressed from several different promoters (see Table 2, Materials and Methods). Most importantly, the expression levels of the Spd-2- and Cnn-fusion proteins used to measure recruitment dynamics were similar to endogenous levels (Figure S1), while the Polo-GFP fusion was expressed from a GFP-insertion into the endogenous Polo gene (Buszczak *et al*, 2007).

**Figure 1.**
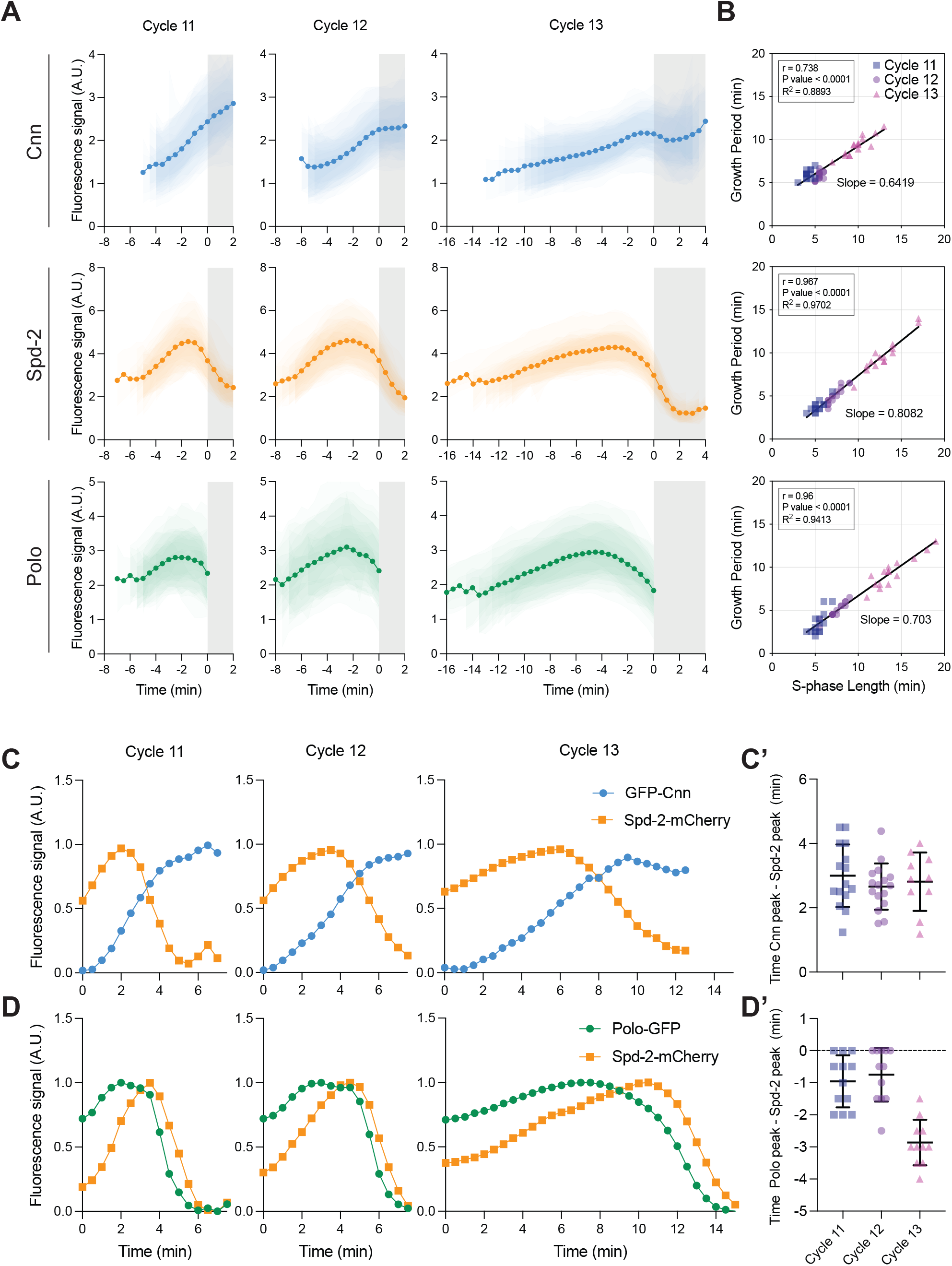
Analysis of PCM scaffold assembly dynamics during nuclear cycles 11-13. **(A)** Graphs show the average centrosomal fluorescence intensity of NG-Cnn, Spd-2-GFP, and Polo-GFP—*dark lines* (±SD for each individual embryo indicated in reduced opacity; N≥15 embryos)—over time during nuclear cycles 11, 12, and 13. The white parts of the graphs indicate S-phase and the grey parts mitosis. All individual embryo tracks were aligned to the start of mitosis (NEB; t=0). **(B)** Scatter plots show the correlation between the centrosome growth period and S-phase length for the embryos analysed in (A). Lines indicate mathematically regressed fits. The goodness of fit (*R^2^*) was assessed in GraphPad Prism. The bivariate Gaussian distribution of the data was confirmed by Henze-Zirkler test, and the strength of correlation (*r*) and the statistical significance (*P-value*) were calculated using Pearson correlation test. **(C, D)** Graphs show the average centrosomal fluorescent intensity over time during nuclear cycles 11, 12 and 13 for embryos (N≥8) co-expressing Spd-2-mCherry (orange) with either GFP-Cnn (blue) (C) or Polo-GFP (green) (D). Fluorescence intensity was rescaled to between 0 and 1 in each cycle. **(C’,D’)** Dot plots compare the time difference between the peak Spd-2-mCherry levels and the peak GFP-Cnn (C’) or peak Polo-GFP (D’) levels in each embryo. Data are presented as Mean±SD.

The rapid nuclear cycles in these embryos comprise alternating periods of S- and M-phase without intervening Gap periods, and S-phase gradually lengthens at each successive cycle (Foe & Alberts, 1983). Perhaps surprisingly, the centrosomal recruitment dynamics of Cnn were quite distinct from Spd-2 and Polo (Figure 1A; Figure S2). In all the nuclear cycles, the centrosomal levels of NG-Cnn increased through most of S-phase, the period when centrosomes grow in preparation for mitosis in these rapidly cycling embryos. In cycle 11, however, NG-Cnn levels continued to increase even after the embryos had entered mitosis—scored by nuclear envelope breakdown (NEB; t=0 in Figure 1A) and indicated by the *grey shading* in the graphs in Figure 1A—while in cycles 12 and 13 centrosomal levels peaked and then largely plateaued at about the time (cycle 12), or a few minutes before (cycle 13), the embryos entered mitosis. In contrast, the centrosomal levels of Spd-2-GFP and Polo-GFP peaked in mid-late S-phase and then started to decline well before NEB (Figure 1A; Figure S2). In these syncytial embryos, S-phase length is determined by the activity of the core Cdk/Cyclin **<u>c</u>**ell **<u>c</u>**ycle **<u>o</u>**scillator (CCO) that drives progression through these early nuclear cycles (Farrell & O’Farrell, 2014; Liu *et al*, 2021), and there was a strong correlation (r~0.96; p<0.0001) between S-phase length and the Spd-2 and Polo growth period (measured as the time it takes for Spd-2 and Polo levels to peak in S-phase) (Figure 1B). This suggests that CCO activity influences the kinetics of centrosomal Polo and Spd-2 recruitment.

Spd-2/Cep192 is thought to be the major protein that recruits Polo into the assembling mitotic PCM in vertebrates (Joukov *et al*, 2014, 2010; Meng *et al*, 2015) worms (Decker *et al*, 2011) and flies (Alvarez-Rodrigo *et al*, 2019), but the shapes of the Spd-2 and Polo centrosomal growth curves were quite distinct, particularly during cycles 11 and 12 (Figure 1A). Moreover, we noticed that during each cycle centrosomal Polo levels peaked slightly before Spd-2 levels peaked, and the centrosomal levels of both Polo and Spd-2 peaked before the levels of Cnn peaked—meaning that the Cnn scaffold could continue to grow and/or plateau even as the centrosomal levels of Polo and Spd-2 declined (Figure 1A). As these measurements were taken from different sets of embryos expressing each protein individually, we confirmed these relative timings in embryos co-expressing Spd-2-mCherry with either Polo-GFP or GFP-Cnn (Figure 1C,D).

### An underlying pulse of Polo activity could explain the observed kinetics of PCM scaffold assembly

As the rise and fall in centrosomal Polo levels appeared to precede the rise and fall in centrosomal Spd-2 levels (Figure 1D), we wondered whether the centrosomes might generate a pulse of Polo activity to initiate the assembly of the Spd-2/Cnn scaffold. We have previously developed a molecular model to explain how Spd-2, Polo and Cnn cooperate to assemble a mitotic PCM scaffold in *Drosophila* embryos (Figure 2A). In this scheme, Spd-2 and Polo are recruited to centrioles, and Spd-2 becomes phosphorylated at centrioles as cells prepare to enter mitosis—allowing Spd-2 to form a scaffold that fluxes outwards away from the centriole (Conduit *et al*, 2014b). This scaffold is structurally weak, but it can bind Polo and Cnn from the cytoplasm, which stabilises the scaffold (indicated by the *dotted line* in Figure 2A). This pool of Polo can then phosphorylate the Cnn to generate an independent Cnn scaffold which is structurally strong and can flux outwards from the Spd-2 scaffold along the centrosomal MTs (Conduit *et al*, 2014a; Feng *et al*, 2017) (see Figure S3 for a cartoon illustration of this scheme).

**Figure 2.**
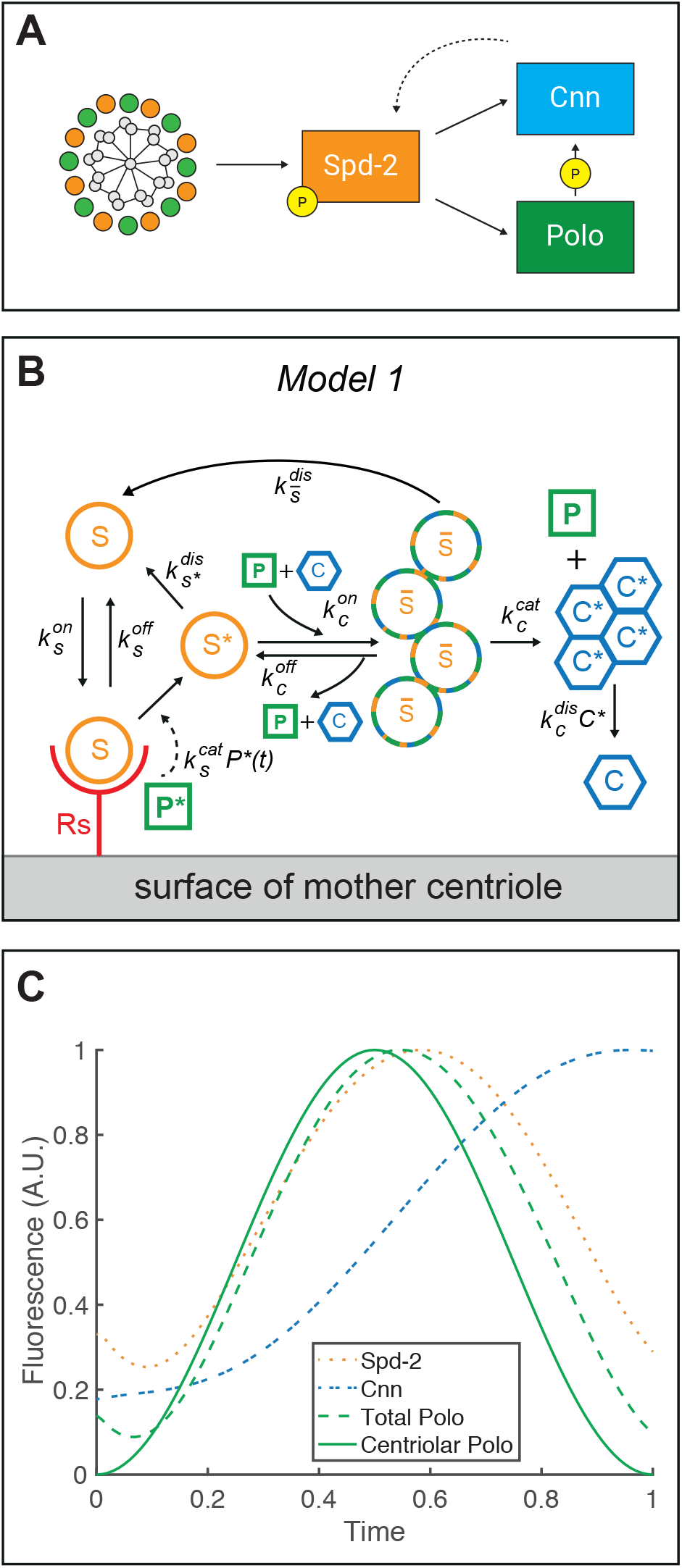
Mathematical modelling of PCM scaffold assembly. **(A)** A schematic summary of the putative molecular interactions that drive the assembly of a Spd-2/Polo/Cnn mitotic PCM scaffold in *Drosophila* (see main text for details). **(B)** Schematic illustrates a version of the molecular model of PCM scaffold assembly that can be formulated as a series of ODEs (see Materials and Methods), allowing us to calculate how the levels of each component in the system changes over time. See main text for the meaning of the various symbols. **(C)** Graph shows the output from the model depicted in (B), illustrating how the centrosomal levels of the various PCM scaffold components change over time if a centriolar pulse of Polo activity (*solid green line*) is imposed on the system. Total Polo (*dotted green line*) represents the sum of the *P*\* generated at the centriole surface and the *P*\* bound to the 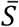 scaffold; Total Spd-2 (*dotted orange line*) represents the sum of Spd-2 in *S*\* and 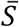; Total Cnn (*dotted blue line*) represents the sum of Cnn in 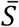 and *C*\*. To better reflect the situation *in vivo*—where the centrosomes start each cycle already associated with some PCM scaffold acquired from the previous cycle (Conduit *et al*, 2010)—we allow the model to run for a complete initial cycle (where the levels of all scaffolding components start at zero) and then graph the behaviour of the system starting from this point during a second cycle. Thus, the pulse of centriolar Polo activity starts from zero at the start of the cycle, but some Polo, Spd-2 and Cnn recruited in the previous cycle are already present at the centrosome.

We turned to mathematical modelling to test whether imposing an underlying pulse of centriolar Polo activity on these proposed molecular interactions could explain the observed kinetics of PCM scaffold assembly. In this model (*Model* 1; Figure 2B) we assume that a pulse of active Polo (*P*\*) is generated at the surface of mother centrioles, with levels peaking at mid-S-phase (we explore below how this pulse might be generated). We allow centrosomal receptors (*R_S_*) to recruit cytoplasmic Spd-2 (*S*) to the centriole to form the complex 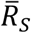. The Spd-2 bound to this complex can be phosphorylated by *P*\* and converted to a form that can form a scaffold (*S*\*) that is released from 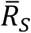 to flux outwards. This scaffold is unstable and can be rapidly converted back to *S* by a phosphatase, which we allow to be active in the cytoplasm at a constant level. However, *S*\* can also bind cytoplasmic Polo (*P*) and Cnn (*C*), to form a more stable scaffold 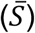 that converts back to *S* relatively slowly. When bound to 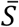, Polo is activated so that it can phosphorylate the 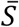-bound Cnn and convert it into a form (*C*\*) that can form a scaffold and be released from 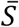 to flux further outwards. In this way, the Spd-2 scaffold acts to convert catalytically *C* into the scaffold *C*\*. The *C*\* scaffold disassembles when it is dephosphorylated by a cytoplasmic phosphatase (PPTase), which we allow to be active in the cytoplasm at a constant level. Note that this PPTase activity drives a low-level of Cnn scaffold disassembly during the assembly process, but it is not intended to mimic the high levels of PPTase activity that are thought to drive the rapid disassembly of the PCM scaffold at the end of mitosis (Enos *et al*, 2018; Magescas *et al*, 2019; Mittasch *et al*, 2020). As explained above, this rapid disassembly is a complex process that we do not attempt to measure or model here. We also allow the rate of *C*\* disassembly to increase as the size of the *C*\* scaffold increases, which appears to be the case in these embryos (see Materials and Methods).

We modelled these reactions as a system of ordinary differential equations (ODEs, detailed in Materials and Methods) and estimated values for each of the 12 model parameters (see “Justification of Model Parameters” in Materials and Methods). Encouragingly, the output of this model recapitulated two of the most surprising features of scaffold assembly dynamics that we observed *in vivo* (Figure 2C): (1) The imposed centriolar *P*\* pulse (*solid green line*, Figure 2B) generated a subsequent pulse in centrosomal Spd-2 levels (*dotted orange line*, Figure 2C); (2) the system generated the assembly of a Cnn scaffold (*dotted blue line*, Figure 2C) that could continue to grow and then plateau even as centrosomal Polo and Spd-2 levels declined. To assess the robustness of this model we tested the effect of individually halving or doubling each of the reaction rate parameters. Although the precise shapes of the curves varied, these two key features were recapitulated in all cases (Figure S4). Thus, this simple model can robustly explain the basic dynamic features of PCM scaffold assembly kinetics that we observe *in vivo* in the parameter regime we consider.

### Spd-2 and Ana1 help to generate the centrosomal Polo pulse

How might the centrioles generate a pulse of Polo activity? This pulse of activity is unlikely to simply reflect the general activity of Polo in the embryo, which, like Cdk/Cyclin activity (Deneke *et al*, 2016), peaks during mitosis (Stefano Di Talia, Duke University (USA), *personal communication*). Thus, the centrioles must generate a local pulse of Polo activity well before Polo is maximally activated in the rest of the embryo more generally. Polo/PLK1 is known to be recruited to mitotic centrosomes by its Polo-box domain (PBD) that binds to phosphorylated S-S(P)/T(P) motifs (Elia *et al*, 2003; Reynolds & Ohkura, 2003; Song *et al*, 2000; Seong *et al*, 2002); this recruitment is sufficient to at least partially activate the kinase (Xu *et al*, 2013). In fly embryos, the Polo required for mitotic PCM assembly appears to be recruited to centrosomes via the sequential phosphorylation of S-S(P)/T(P) motifs first in Ana1 (that recruits Polo to mother centrioles) (Alvarez-Rodrigo *et al*, 2021) and then in Spd-2 (which recruits Polo into the expanding mitotic PCM) (Alvarez-Rodrigo *et al*, 2019).

To test the potential role of these proteins in generating the Polo pulse we examined Polo-GFP recruitment during nuclear cycle 12 in embryos expressing a mutant form of either Ana1 (Ana1-S34T-mCherry) (Alvarez-Rodrigo *et al*, 2021) or Spd-2 (Spd-2-S16T-mCherry—previously called Spd-2-CONS-mCherry) (Alvarez-Rodrigo *et al*, 2019) in which multiple S-S/T motifs (34 for Ana1 and 16 for Spd-2) were mutated to T-S/T (Figure 3). These conservative substitutions severely impair the ability of the mutant proteins to recruit Polo, seemingly without perturbing other aspects of their function (Alvarez-Rodrigo *et al*, 2019, 2021). These experiments were performed in the presence of endogenous, untagged, Spd-2 or Ana1 because embryos laid by females co-expressing Polo-GFP in the presence of only Ana1-S34T or Spd-2-S16T die very early in development due to centrosome defects (Alvarez-Rodrigo *et al*, 2019, 2021)—as centrosomes are essential for early embryo development (Stevens *et al*, 2007; Varmark *et al*, 2007), but not for the rest of development in *Drosophila* (Basto *et al*, 2006).

**Figure 3.**
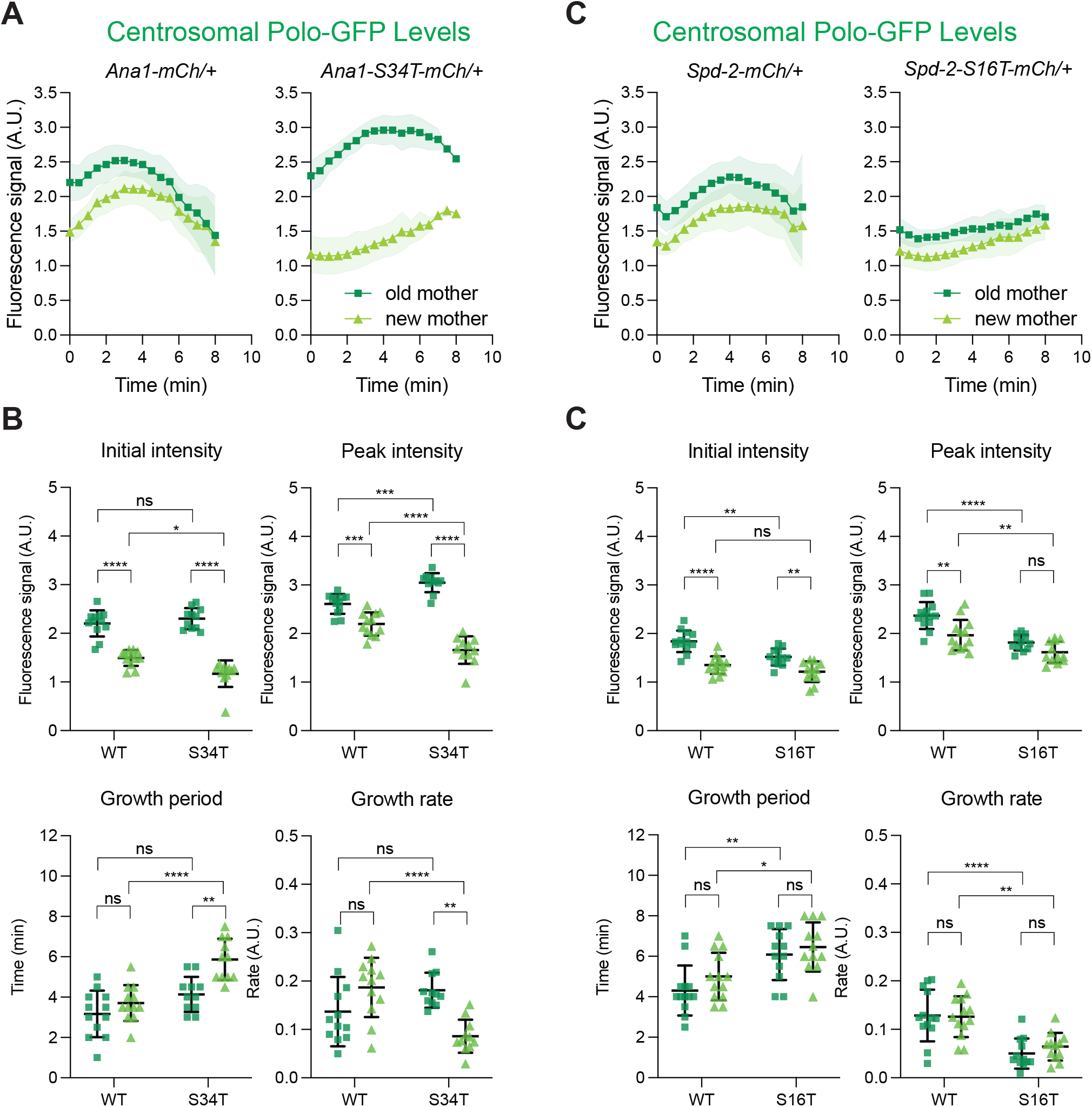
Perturbing the ability of Ana1 or Spd-2 to recruit Polo perturbs the pulse of Polo activity. **(A)** Graphs show how the average fluorescent intensity (±SD) of Polo-GFP changes over time at OM (*dark green squares*) or NM (*light green triangles*) centrosomes during nuclear cycle 12 in embryos (N=12) laid by WT females expressing either Ana1-mCherry, Ana1-S34T-mCherry, Spd-2-mCherry, or Spd-2-S16T-mCherry. In this experiment, embryos were aligned to the start of S-phase (t=0), which was scored by centriole separation. **(B, C)** Bar charts compare various growth parameters (indicated above each graph) of the embryos analysed in (A); dots representing the behaviour of OM and NM centrosomes in each class of embryo are shown in *dark green squares and light green triangles*, respectively. Statistical significance was first assessed by an ordinary two-way ANOVA, and then a Šídák’s multiple comparisons test (*: P<0.05, **: P<0.01, ***: P<0.001, ****: P<0.0001, ns: not significant).

In these experiments, we examined the centrosomes organised by the old-mother centriole (hereafter OM centrosomes) and new-mother centriole (hereafter NM centrosomes) separately, as they behaved differently. In embryos expressing Polo-GFP and WT-Ana1-mCherry, the Polo pulse was similar on OM and NM centrosomes, although NM centrosomes initially organised significantly less Polo than OM centrosomes (left graph, Figure 3A). This is because at the start of S-phase NMs are recruiting Polo for the first time, whereas OMs retain some of the mitotic PCM that they had recruited in the previous cycle (Conduit *et al*, 2010). Although this asymmetry is essentially eliminated by the end of S-phase in normal embryos, it is present at the start of each nuclear cycle because both the OM and NM centrioles become OMs in the *next* cycle; all of the NMs in a cycle are derived from the daughters generated in the previous cycle, and these centrioles do not start to recruit Polo until they mature into mothers during mitosis (Novak *et al*, 2016).

In embryos expressing Polo-GFP and Ana1-S34T-mCherry, the Polo pulse was relatively normal on OMs although, surprisingly, the amount of Polo recruited to OMs increased significantly (right graph, Figure 3A; quantification shown in Figure 3B) (possible reasons for this are discussed in the last Results Section). The Polo pulse was more dramatically perturbed on NMs, exhibiting a reduced growth rate, a lower amplitude and a longer period (Figure 3A,B). We believe that Ana1-S34T more dramatically perturbs NM centrosomes because Ana1 is only required to recruit Polo to the centrioles (and not to the PCM). We showed previously that once some mitotic PCM has been established around a centriole (as is the case at OM centrosomes), it can help recruit Polo to centrosomes and so partially bypass the requirement for Ana1 to initiate Polo recruitment to the centrioles; thus, Ana1 is more important at NMs, which are recruiting Polo for the first time (Alvarez-Rodrigo *et al*, 2021).

The Polo-GFP pulse was relatively normal on OM and NM centrosomes in embryos co-expressing WT Spd-2-mCherry (right graph, Figure 3C), but was dramatically perturbed at both centrosomes in embryos expressing Spd-2-S16T-mCherry (left graph, Figure 3C)—exhibiting a reduced growth rate, a lower amplitude and a longer period (Figure 3D). We believe that Spd-2-S16T-mCherry effects both centrosomes equally because Spd-2 is primarily responsible for recruiting Polo-GFP to the mitotic PCM (rather than to the centrioles), so OM centrosomes cannot establish a relatively normal mitotic PCM in the presence of Spd-2-S16T, which they can do in the presence of Ana1-S34T. Taken together, these results indicate that Ana1 and Spd-2 play an important part in generating the centrosomal Polo pulse, with Ana1 having the dominant role in initially recruiting Polo to the centrioles and Spd-2 having the dominant role in subsequently recruiting Polo to the expanding mitotic PCM.

### Mathematical modelling indicates that an interaction between Ana1 and Polo could generate a centrosomal pulse of Polo activity

Intriguingly, expressing either Ana1-S34T or Spd-2-S16T in embryos perturbed not only the amplitude of the Polo pulse, but also its period (Figure 3). Moreover, in embryos expressing Ana1-S34T the period was significantly perturbed on NM centrosomes but not on OM centrosomes—even though these centrosomes are located very close to each other in the same cytoplasm. This suggests that the kinetics of the Polo pulse are generated by mechanisms that act locally on individual centrosomes, rather than globally on the embryo as a whole. These observations also suggest that Polo itself might ultimately inhibit its own recruitment to centrosomes, as in Ana1-S34T embryos OM centrosomes (with high levels of Polo) stop recruiting Polo before NM centrosomes (that have lower levels of Polo) (Figure 3A,B). With this in mind, we developed a simple mathematical model to explain how an interaction between Polo (*P*) and its centriolar Receptor (*R_P_*) (in these embryos most likely Ana1) could generate a pulse of Polo activity.

In this model (*Model 2*; Figure 4A), the centriolar Polo receptor is initially in an inactive state that cannot recruit Polo 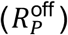. Mitotic PCM recruitment is initiated when 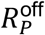 is phosphorylated on S-S(P)/T(P) motifs by a kinase (whose activity is potentially regulated by the CCO, see below) to generate *R_P_*. These activated receptors can recruit and activate cytoplasmic Polo to form the complex 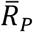. This pool of centriole-bound active Polo can phosphorylate the Spd-2 bound to the centriolar Receptor complex 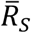—also potentially Ana1/CEP295 (Tsuchiya *et al*, 2016)—to generate *S*\*. This initiates mitotic PCM scaffold assembly (as described in *Model 1*; Figure 2A). Crucially, we also allow the centriole-bound active Polo to slowly phosphorylate 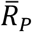 at additional sites (i.e. not the original S-S/T motifs) to generate 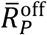, which can no longer recruit Polo. In this way, an activator (*R_P_*), activates its own inhibitor (*P*\*) to form a classical delayed negative feedback network (Novák & Tyson, 2008) to generate a local pulse of Polo activity at the centriole (*solid green* line, Figure 4B). We speculate that this system is reset when 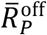 is dephosphorylated during mitosis by a phosphatase to regenerate 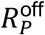 (this step is not depicted in the schematic and we do not model it here). When we used the pulse of Polo activity generated by *Model 2* to feed into *Model 1* to generate the PCM scaffold, it produced assembly kinetics that were similar to the original *Model 1* (where we simply imposed a Polo pulse on the system) (Figure 4B). Hence, *Model 2* can plausibly explain how centrioles might generate a pulse of Polo activity.

**Figure 4.**
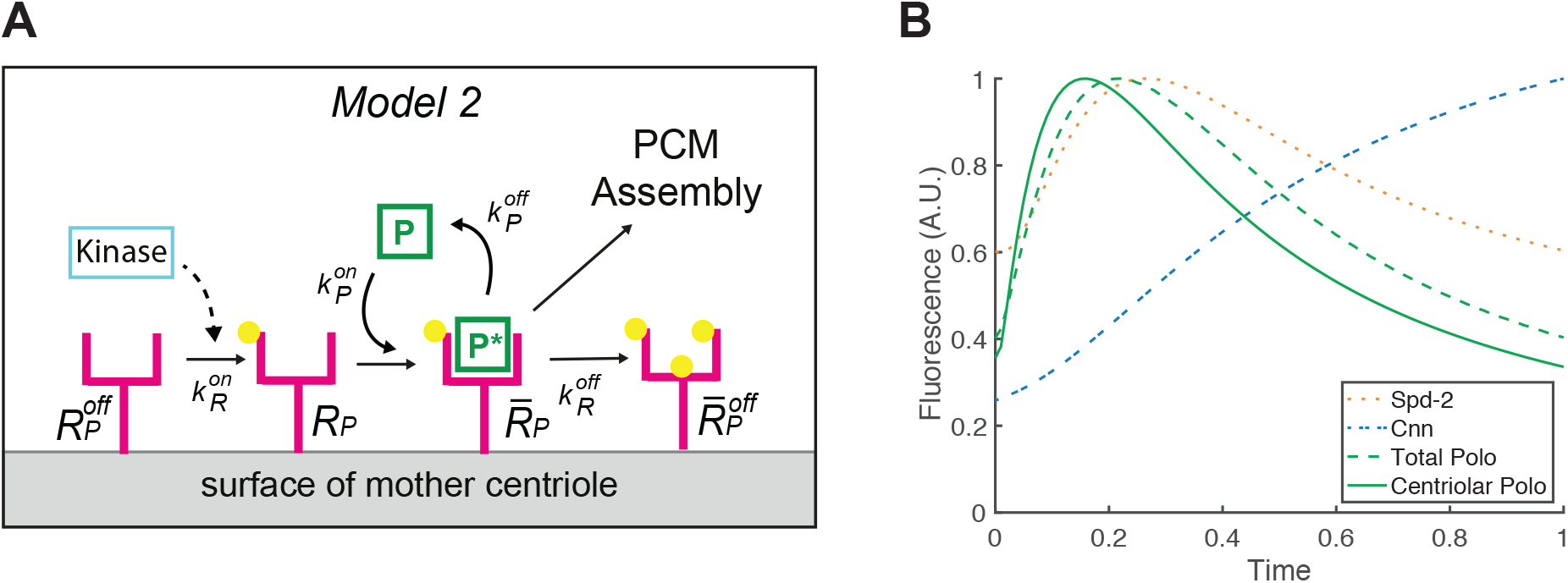
Mathematical modelling can explain how centrioles might generate a pulse of Polo activity. **(A)** Schematic illustrates a molecular model (*Model 2*) of how an interaction between Polo and its centriolar receptor can generate an oscillation in centriolar Polo levels (see main text for details). **(B)** This model was formulated as a series of ODEs (see Materials and Methods) to allow us to graph how the levels of centriolar Polo would change over time (*solid green line*). The graph also illustrates the output when this pulse of Polo activity is fed into our earlier model of PCM scaffold assembly (Model 1, Figure 2A)—illustrating how the levels of total centrosomal Polo (*dotted green line*), Spd-2 (*dotted orange line*) and Cnn (*dotted blue line*) (as defined in the legend to Figure 2) change over time. As in Figure 2B, we allow the model to run for a complete initial cycle and then graph the behaviour of the system during a second cycle.

As Polo/PLK1 turns over rapidly at centrosomes (Kishi *et al*, 2009; Mahen *et al*, 2011) (see below), it seems likely that the centrosomal Polo receptors (likely Ana1 and Spd-2) constantly generate and then release active Polo, which may have some ability to diffuse and phosphorylate local targets before it is inactivated. This is not considered in our simple model, but it would explain why expressing either Ana1-S34T or Spd-2-S16T lengthens the period of the Polo pulse (Figure 3). If the centriole and PCM receptors (Ana1 and Spd-2, respectively) recruit less Polo, the centriole receptor (Ana1) will be inactivated more slowly. Thus, in the presence of Spd-2-S16T or Ana1-S34T, Polo would be recruited more slowly, but for a longer period—as we observe (Figure 3).

### The rate of Polo recruitment and PCM scaffold growth is influenced by the Cdk/Cyclin cell cycle oscillator (CCO)

The period of centrosomal Polo recruitment is strongly correlated with S-phase length (Figure 1B), which is determined by CCO activity (Farrell & O’Farrell, 2014; Liu *et al*, 2021). We speculate that in our molecular model the CCO could influence Polo recruitment by regulating the rate 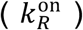 at which the relevant protein kinase phosphorylates the centriolar Polo receptor (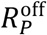, likely Ana1) (Figure 4A). If this receptor is initially phosphorylated more slowly (if the CCO is less active), then Polo will be recruited more slowly and the PCM scaffold will grow more slowly. During nuclear cycles 10-13, the rate at which the CCO is activated during S-phase naturally slows at successive cycles (Farrell & O’Farrell, 2014; Liu *et al*, 2021). Thus, if our model is correct, the rate of Polo recruitment to the centrosome should slow at each successive nuclear cycle, and the rate of Spd-2 and Cnn scaffold growth should also slow. To test if this was the case we performed a Fluorescence Recovery After Photobleaching (FRAP) analysis (Figure 5; Figure S5). The fluorescent signal of the three PCM scaffold proteins recovered at very different rates (note the different timescales in Figure 5A-C) with Polo turning over with a *t_1/2_* of ~10secs and Spd-2 and Cnn fluorescence recovering more slowly. Strikingly, however, the average rate of fluorescence recovery of all three proteins slowed at successive cycles (Figure 5B), consistent with our hypothesis that these parameters are influenced by the CCO.

**Figure 5.**
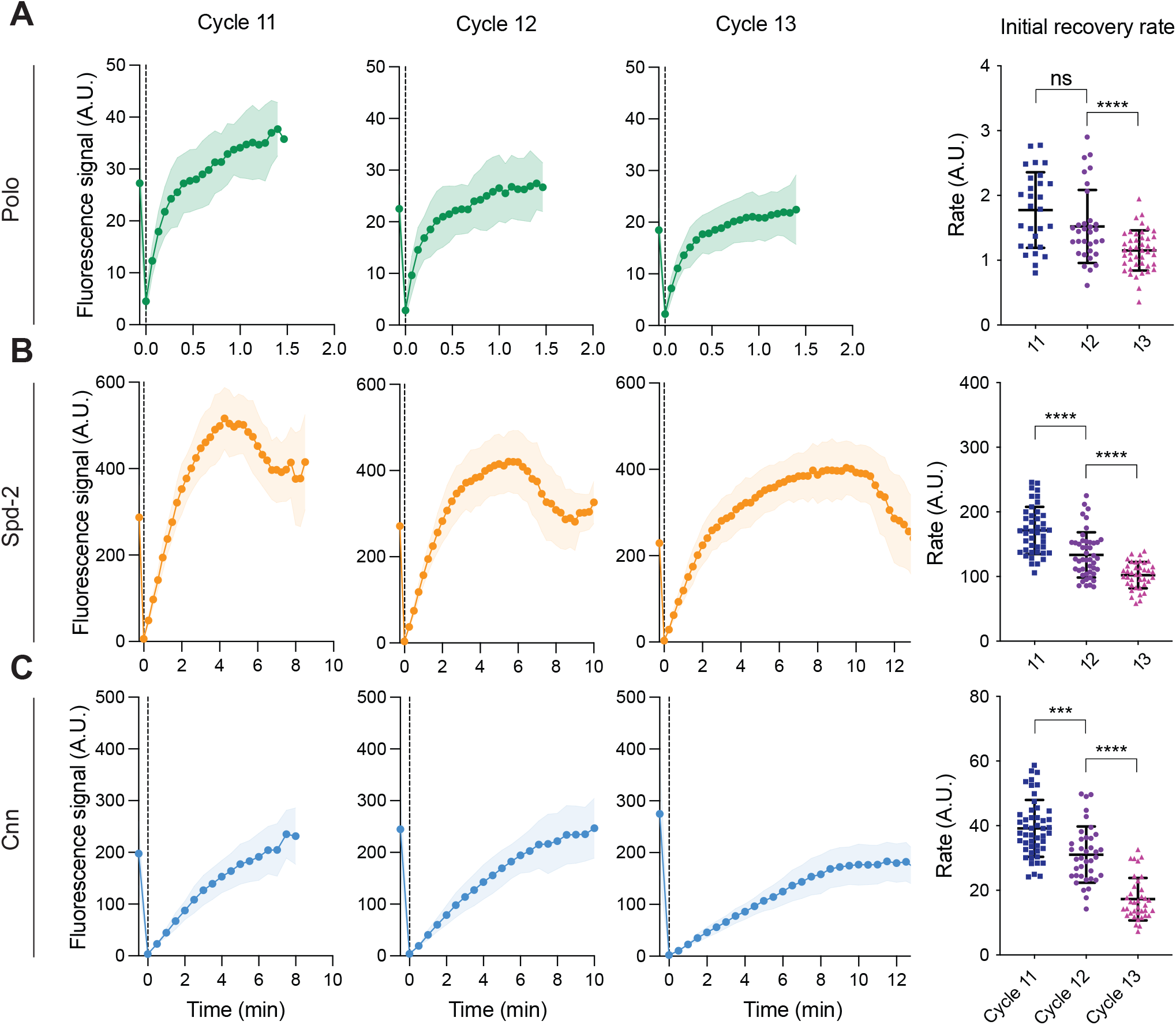
The rate of recruitment of Polo, Spd-2 and Cnn slows during successive nuclear cycles. Graphs show the recovery of centrosomal fluorescence intensity (±SD) after photobleaching of Polo-GFP (A; imaged every 4secs), Spd-2-NG (B; imaged every 15secs) and NG-Cnn (C; imaged every 30secs) in early S-phase of cycle 11, 12 or 13. Each coloured line is an average of a total of 25-50 centrosomes imaged from 10-12 embryos in each cycle. Dot plots compare the average initial recovery rate (±SD) of these proteins in nuclear cycle 11, 12, or 13. Statistical significance was assessed using a Dunnett’s T3 multiple comparison test (ns: not significant, ***: P<0.001, ****: P<0.0001) after performing a Welch ANOVA test.

In interpreting this experiment it is important to remember that Spd-2 and Cnn do not “turn-over” at centrosomes in the classical sense, as both proteins incorporate into the PCM in the central region around the mother centriole and then flux outwards to leave the PCM from the more peripheral regions (Conduit *et al*, 2010, 2014b). Thus, the initial rate of fluorescence “recovery” that we measure for Spd-2 and Cnn largely reflects the growth of the scaffold (i.e. the incorporation of new Spd-2 and Cnn molecules into the central region) rather than the turn-over of molecules already incorporated into the PCM. This is not the case for Polo, which turns-over rapidly throughout the entire PCM volume as it binds and unbinds from its centrosomal receptors (Conduit *et al*, 2014b). Thus, we believe the decreased rate of Polo turnover at centrosomes at successive cycles reflects the slowing rate at which its receptors are activated by the CCO at successive cycles, while the decreasing rate of Spd-2 and Cnn addition at successive cycles reflects the slower rate of scaffold growth caused by the slower recruitment of Polo.

### Mathematical modelling predicts the consequence of lowering the concentration of either Spd-2 or Ana1

To test whether our mathematical models have predictive power, we used them to model the consequences of halving the amount of either Spd-2 or Ana1 in the system, while we measured experimentally the Polo-GFP pulse in embryos laid by females expressing only one copy of *Spd-2* (*Spd-2*^+/-^ embryos) or *ana1* (*ana1*^+/-^ embryos) (Figure 6). Although the precise shape of the growth curves generated by the mathematical model (Figure 6A,C) did not match closely the experimental data (Figure 6B,D) (potential reasons for this are discussed below), the model correctly predicted that halving Spd-2 levels would lead to a general reduction in centrosomal Polo recruitment (Figure 6A). Perhaps more convincingly, the model also correctly predicted the surprising observation that although halving Ana1 levels led to an initial reduction in centrosomal Polo levels (as one might intuitively predict), the centrosomes in the *ana1*^+/-^ embryos ultimately associated with *more* Polo than controls by the end of S-phase (Figure 6B). In our model, this occurs because the peak of the Polo pulse is shifted to later in S-phase and its rate of decline is shallower in the half-dose *ana1*^+/-^ embryos—because the Ana1-Polo receptors are inactivated more slowly.

**Figure 6.**
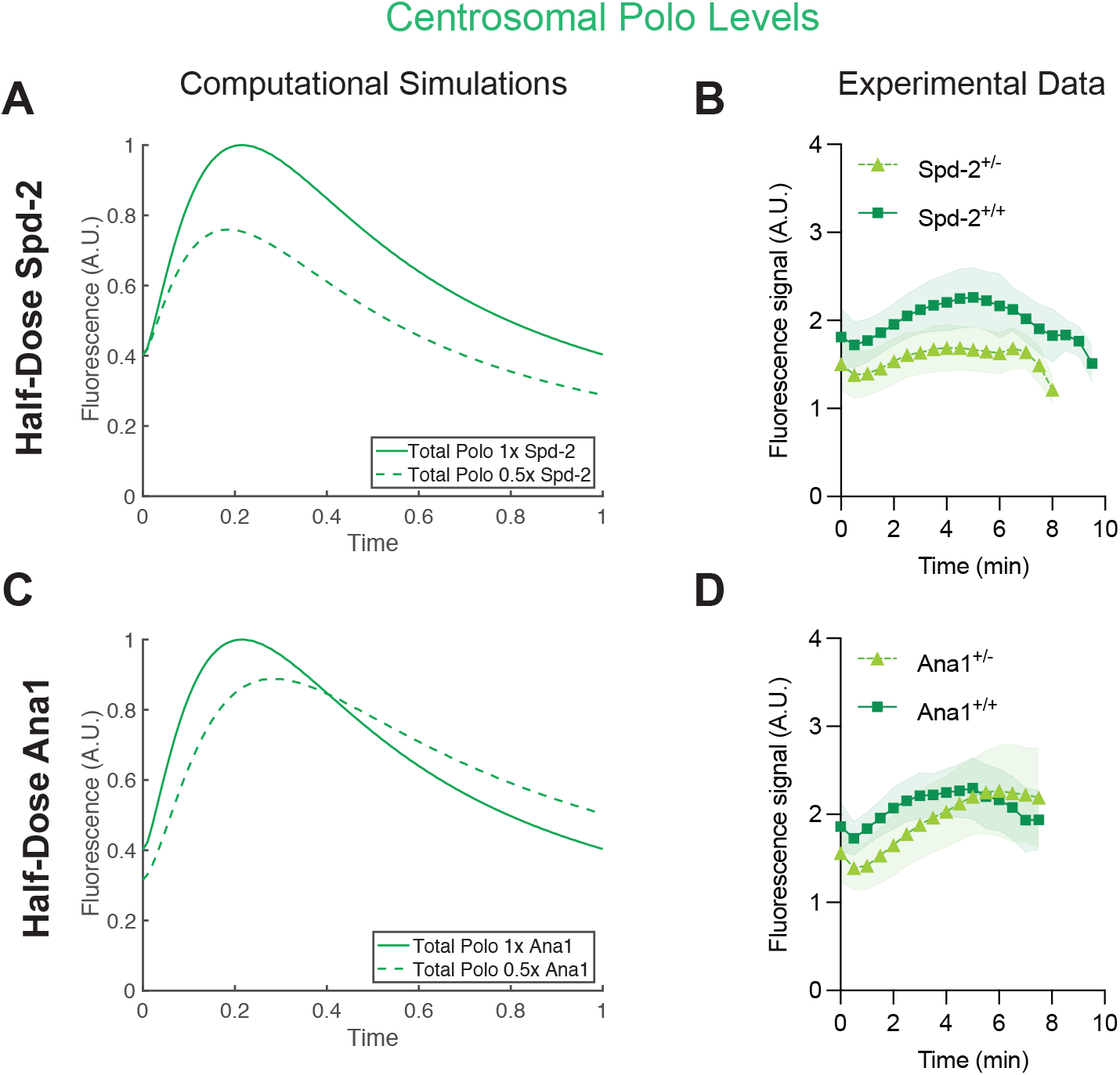
Mathematical models can predict the broad behaviour of the Polo pulse when the genetic dosage of *Spd-2* or *ana1* is halved. (**A,B**) Graphs compare mathematical predictions of the Polo pulse (generated using Model 2) in embryos expressing normal levels of Spd-2 (A) and Ana1 (B) (*solid green lines*) or in embryos where the levels of Spd-2 (A) or Ana1 (B) in the system have been halved (*dotted green lines*). (**C,D**) Graphs show *in vivo* data showing how the average centrosomal fluorescent intensity (±SD) of Polo-GFP changes over time during nuclear cycle 12 in embryos (N≥12) laid by either WT (+/+) females (*dark green squares*), or females in which the genetic dosage of *Spd-2* (A) or *ana1* (B) has been halved (+/-) (*light green triangles*).

To test the robustness of these predictions, we performed a “parameter sweep” (Figure 7), individually varying each of the 13 reaction rate parameters between 0.5X and 2X its initial value in either WT and half-dose Ana1 conditions (Figure 7, top graphs) or WT and half-dose Spd-2 conditions (Figure 7, bottom graphs). We monitoring how these changes in parameter values affected the relative behaviour of the centrosomal Polo pulse in the WT and half-dose embryos in terms of: (1) The peak amount of Polo recruited (Figure 7A); (2) The timing of the Polo peak (Figure 7B); (3) The final amount of Polo recruited at the end of S-phase (Figure 7C). This analysis revealed that under certain parameter conditions the peak amount of Polo recruited to centrosomes was greater in the 0.5X Ana1 embryos than the WT Ana1 embryos (so on the graph the ratio of these values is >1, indicated by the *pink boxing*, Figure 7A), whereas under other parameter conditions the peak amount of Polo recruited to centrosomes was greater in the WT embryos (so on the graph the ratio of these values is <1, indicated by the *unboxed white areas*, Figure 7A). In contrast, under all parameter conditions tested, the timing of the centrosomal Polo peak was shifted to later in the cycle in the Ana1 half-dose embryos, but earlier in the cycle in the Spd-2 half dose embryos (Figure 7B), while the amount of Polo recruited to centrosomes at the end of S-phase was larger in the Ana1 half-dose embryos and smaller in the Spd-2 half-dose embryos (Figure 7C).

**Figure 7.**
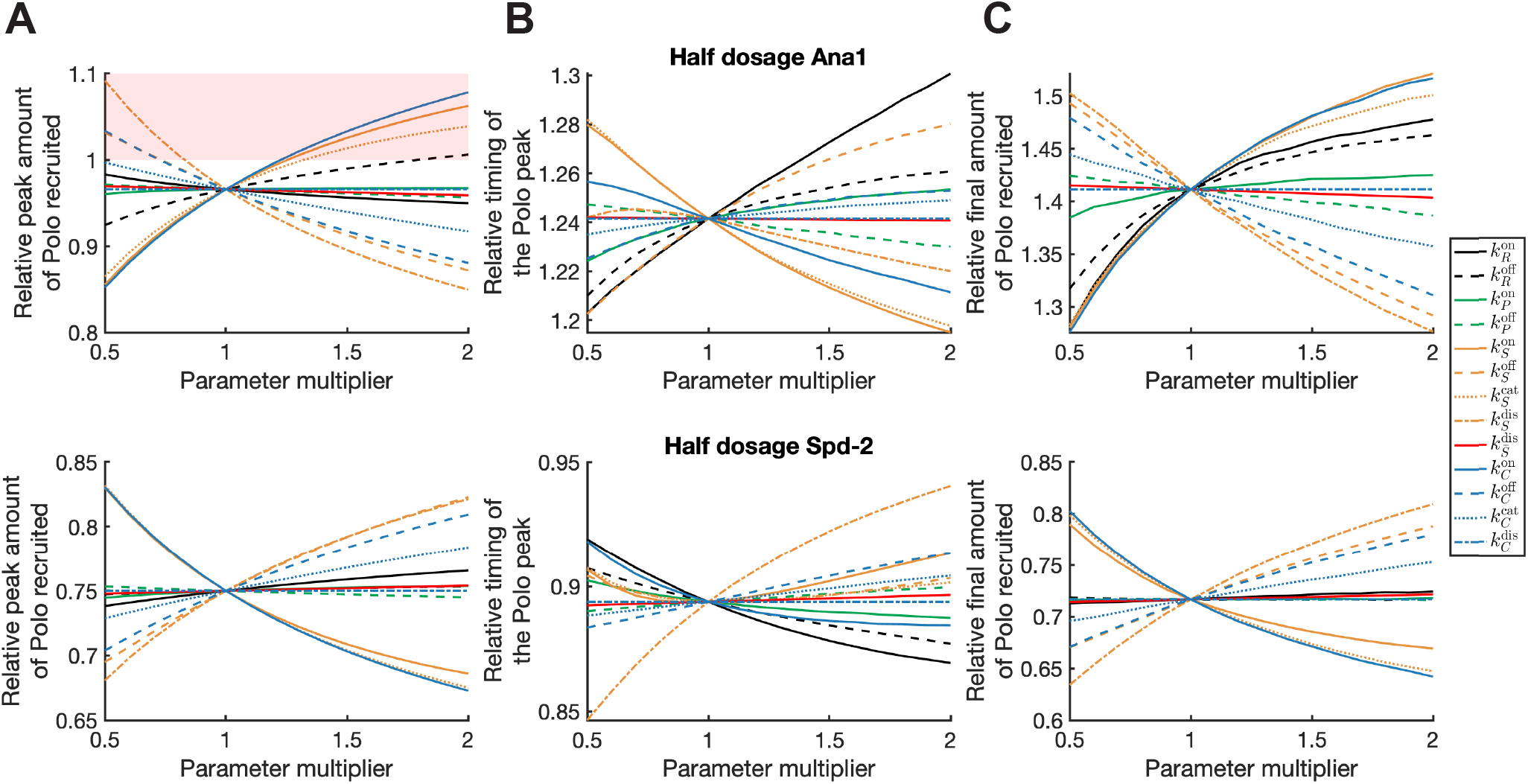
Model predictions of Polo behaviour are robust to changes in parameter values. Graphs show how the relative behaviour of the Polo pulse is predicted to change between WT and either Ana1 (top graphs) or Spd-2 (bottom graphs) half-dose embryos when the 13 reaction rate parameters are individually varied between 0.5-2X of their initial value (which is represented by 1 on the x-axis). Values <1 on the y-axis indicate that the half-dose embryos have a lower value for the corresponding attribute than WT embryos, and values > 1 indicate that the half-dose embryos have a greater value for the corresponding attribute than WT embryos. The behaviours analysed are: **(A)** The relative peak amount of Polo recruited to centrosomes; **(B)** The relative timing of the Polo peak; **(C)** The relative final amount of Polo recruited to centrosomes at the end of S-phase. Conditions in which the model incorrectly predicts the behaviour of the Polo pulse under half-dose conditions are *boxed in pink*. For example, the peak amount of Polo recruited to centrosomes is predicted to be higher in the WT embryos (as is observed experimentally) in most parameter regimes, but under some parameter values it is predicted to be higher in the Ana1 half-dose embryos (A, *pink box*, top graph). By contrast, the surprising experimental observation that the amount of Polo recruited to centrosomes at the end of S-phase is higher in the Ana1 half-dose embryos is predicated under all parameter values tested—indicated by the absence of a pink box (C, top graph). Similarly, the behaviour of all other attributes are correctly predicted in all of the parameter regimes considered.

We conclude that the surprising observation that centrosomal Polo levels are actually higher at the end of S-phase in the Ana1 half-dose embryos is robustly predicted by Model 2. Importantly, this finding could explain our earlier puzzling observation that OM centrosomes recruit more Polo than normal in embryos expressing some Ana1-S34T protein (Figure 3A). If these centrosomes associate with excessive Polo at the end of S-phase, this could lead to the gradual accumulation of Polo at OM centrosomes over multiple division cycles.

Finally, we note that these models are purposefully minimal to reduce the number of parameters and test possible mechanisms rather than to mimic experimental data. This approach explains why the overall shape of the predicted growth curves do not exhibit all of the finer characteristics of the experimental data. For instance, in our models the Polo and Spd-2 pulses consistently have higher amplitudes and earlier peaks compared to experimental data (see, for example, Figure 6). In principle, this problem can be solved by introducing additional intermediate steps into the model (e.g. by requiring that the centriolar receptors are phosphorylated on multiple sites to be activated). Any such additional steps will delay the system (and so shift the peaks to later in S-phase), and also allow the receptors to simultaneously occupy a larger distribution of states (and so dampen the amplitude). Since such intermediate states are likely to exist—Ana1 and Spd-2, for example, both appear to be phosphorylated on more than one site to recruit Polo (Alvarez-Rodrigo *et al*, 2019, 2021)—we acknowledge the consequences of neglecting them in our model but choose to do so in favour of simplicity.

## Discussion

We show here that the mother centrioles in the early *Drosophila* embryo generate a pulse of Polo activity at the start of each nuclear cycle, and we propose that this initiates centrosome maturation by catalysing the local assembly of a Spd-2/Cnn PCM scaffold. In the early *Drosophila* embryo, the global cytoplasmic activity of mitotic kinases such as Cdk1 and Polo are relatively low at the start of each nuclear cycle, but they rise steadily during S-phase to peak during mitosis (Deneke *et al*, 2016) (Stefano Di Talia, personal communication). Thus, the local activation of Polo at centrioles precedes the global rise in embryonic Polo activity that occurs as embryos enter mitosis.

Perhaps surprisingly, centrosomal Polo levels actually start to decrease before mitotic entry in early fly embryos, suggesting that the local activation of Polo at centrosomes may not be required to maintain the mitotic PCM once the embryos have entered mitosis. We speculate that this is because global Polo activity in the embryo is high during mitosis, and this is sufficient to maintain the already assembled mitotic PCM. We envisage that high levels of global Polo (and perhaps other mitotic kinase) activity would not only maintain the phosphorylation of the Cnn scaffold (and perhaps that of other mitotic PCM components), but would likely also suppress the activity of protein phosphatases (PPTases) that promote the disassembly of the mitotic PCM (Glover, 2012). Such PPTase’s have been identified in worm embryos (Enos *et al*, 2018; Magescas *et al*, 2019; Mittasch *et al*, 2020), and are likely to perform a similar function in fly embryos.

Recent experiments in the early *C. elegans* embryo indicate that centriolar Polo/PLK-1 activity is also required to initiate mitotic PCM assembly in this system. Worm embryos build a mitotic PCM scaffold using a PLK-1/SPD-2/SPD-5 system that is analogous to the fly Polo/Spd-2/Cnn system (Hamill *et al*, 2002; Kemp *et al*, 2004; Pelletier *et al*, 2004; Dammermann *et al*, 2004; Woodruff *et al*, 2015a; Laos *et al*, 2015; Wueseke *et al*, 2016; Ohta *et al*, 2021). In these embryos, the centrioles and PLK-1 are both continuously required to promote the *growth* of the mitotic PCM (Cabral *et al*, 2019), but the centrioles are not required to *maintain* the fully grown mitotic PCM—although PLK-1 activity is still essential, and is presumably provided from the cytoplasm (Cabral *et al*, 2019). Thus, as in fly embryos, the centrioles appear to provide a source of Polo/PLK-1 activity that initiates and sustains the growth of the mitotic centrosome prior to mitotic entry (Zwicker *et al*, 2014; Cabral *et al*, 2019), but this centriolar source does not appear to be required once the embryos enter mitosis. We speculate that in worms, as in flies, this is because global PLK-1 activity is sufficiently high during mitosis to maintain the mitotic PCM.

We propose that the ability of centrioles to generate a local activation of Polo prior to mitotic entry may be a universal feature of centrosome maturation, not just a specialisation of embryos. In many cells, the centrosomes start to mature prior to NEB, so presumably well before Polo is fully activated in the cytoplasm. In such systems, mother centrioles could initiate centrosome maturation by locally activating Polo/PLK1 prior to mitosis.

Although high levels of centrosomal Polo/PLK1 may not be required to sustain the mitotic PCM once cells are in mitosis, centrosomal levels of PLK1 nevertheless remain relatively high during mitosis in human cells and worm embryos and only rapidly decline as cells exit mitosis (Golsteyn *et al*, 1995; Mittasch *et al*, 2020). It is therefore unclear why Polo levels decrease prior to the entry into mitosis in *Drosophila* embryos. A possible explanation is that fly embryos cycle extremely rapidly, and mitotic PCM disassembly is initiated at the end of mitosis at essentially the same time as the newly disengaged centrioles start to initiate a new round of mitotic PCM recruitment (Conduit *et al*, 2010). In this scenario, the partial disassembly of the Spd-2/Polo scaffold prior to mitosis might be important to allow the pre-existing mitotic PCM to disassemble efficiently at the end of mitosis, while at the same time Ana1 is starting to recruit Polo to centrioles to initiate a new round of mitotic PCM assembly. Interestingly, as in fly embryos, PLK-1 also appears to be the first mitotic PCM component to leave the centrosome in worm embryos—although it only does so at the end of mitosis (Mittasch *et al*, 2020). Thus, although the details and precise timing will vary, the ability of centrioles to switch-on, and then switch-off, the local activation of Polo/PLK1 may be a common feature of centrosome maturation.

Our data and mathematical modelling are consistent with the possibility that Polo is recruited and activated at centrioles to initiate mitotic PCM assembly by its interactions with phosphorylated S-S(P)/T(P) motifs in Ana1 and Spd-2. Although PLK1 binding to these motifs can activate its kinase activity (Xu *et al*, 2013), other centrosomal kinases, such as Cdk1/cyclin B and Aurora A, may also be required to fully activate Polo/PLK1 at the centrioles/centrosomes. Spd-2/CEP192 proteins appear to be universally required for mitotic centrosome assembly (Kemp *et al*, 2004; Pelletier *et al*, 2004; Gomez-Ferreria *et al*, 2007; Zhu *et al*, 2008; Dix & Raff, 2007; Giansanti *et al*, 2008), and there is strong evidence that they recruit Polo/PLK1 (Decker *et al*, 2011; Joukov *et al*, 2014; Meng *et al*, 2015; Alvarez-Rodrigo *et al*, 2019) and also Aurora A to promote their mutual activation (Joukov *et al*, 2014). In both flies and humans Ana1/CEP295 proteins are required for centrosome maturation (Izquierdo *et al*, 2014; Fu *et al*, 2016; Tsuchiya *et al*, 2016; Saurya *et al*, 2016). Although *C.elegans* do not have an obvious Ana1 homologue, proteins such as SAS-7 (Sugioka *et al*, 2017) or PCMD-1 (Stenzel *et al*, 2021) could perform an analogous function. Thus, it seems likely that at least elements of the fly system that generates the pulse of centriolar Polo activity will be conserved.

Finally, our studies reveal intriguing similarities between the proposed mechanisms that regulate the growth of the daughter centriole (Aydogan *et al*, 2018, 2020) and the growth of the mitotic PCM. In both cases, centrioles induce a local pulse in the activity of a key enzyme (Plk4 or Polo/PLK1) that regulates the incorporation of key building blocks (Ana2/Sas-6 or Spd-2/Cnn) into an organelle scaffolding structure (the centriole cartwheel or the mitotic PCM scaffold). Moreover, both systems are normally entrained in the embryo by the core Cdk/Cyclin cell cycle oscillator (CCO) to ensure that organelle assembly not only occurs in the right place, but also at the right time. Could similar principles regulate the growth of other organelles? It is becoming increasingly clear that the biogenesis of several membrane bound organelles occurs primarily at specialised contact sites where key activities are concentrated (Wu *et al*, 2018; Farré *et al*, 2019; Prinz *et al*, 2020). It will be interesting to determine if these key activities are recruited to these sites in a pulsatile fashion, and, if so, whether these activity pulses can be entrained by master oscillators such as the CCO and/or the circadian clock.

## Acknowledgements

We are grateful to Stefano di Thalia (Duke University, USA) and Thomas Steinacker (Raff Lab) for communicating unpublished data, to Lisa Gartenmann for generating the Ubq-NG-Cnn line used in this study, and to Alan Wainman for help with microscopy as part of the Micron Oxford Advanced Bioimaging Unit—partly funded by a Strategic Award from the Wellcome Trust (107457). We thank members of the Raff Laboratory for advice, discussion and for critically reading the manuscript. The research was funded by a Wellcome Trust Senior Investigator Award (215523) to J.W.R., an EPSRC award (EP/R020205/1) and a John Fell Fund Award to A.G., and a CRUK Oxford Centre Prize DPhil Studentship (C5255/A23225), a Balliol Jason Hu Scholarship and a Clarendon Scholarship (to S.S.W.).

## Author contributions

This study was conceptualised by S.S.W., Z.M.W., I. A-R., A.G., and J.W.R. Investigation was done by S.S.W. and Z.M.W. Key reagents were generated by S.S. Computational analysis pipelines were developed by S.S.W., K.Y.C. and F.Y.Z. Data was analysed by S.S.W., Z.M.W., A.G., and J.W.R. The project was supervised and administered by A.G. and J.W.R. The manuscript was initially drafted by S.S.W., Z.M.W., A.G., and J.W.R. and all authors contributed to the editing of the manuscript.

## Supplementary Figure Legends

**Figure S1.**
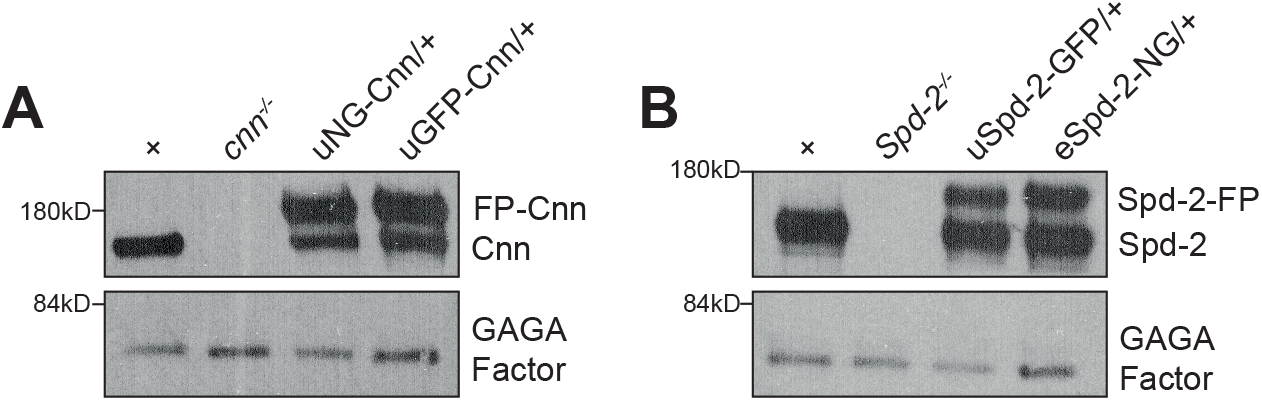
Analysing the relative expression levels of the fluorescent fusion-proteins used to quantify centrosome recruitment dynamics. **(A,B)** Western blots compare the relative expression levels in syncytial embryos of the GFP- and NG-fusion proteins (FP) used here to quantify the centrosomal recruitment dynamics of Cnn (A) or Spd-2 (B). This analysis reveals that GFP-Cnn and NG-Cnn expressed transgenically from the ubiquitin promoter (u) are present at slightly higher levels than the endogenous untagged Cnn. In contrast, Spd-2-GFP expressed transgenically from the ubiquitin promoter (u) and Spd-2-NG expressed as a CRISPR knock-in at the endogenous Spd-2 locus (e) are both present at slightly lower levels than the endogenous protein. Western blots of serial-dilutions of these samples indicate that the uNG-Cnn and uGFP-Cnn are overexpressed by ~2-3 fold compared to the endogenous protein, and that uSpd-2-GFP and eSpd-2-NG are underexpressed by ~2 fold. These blots were also probed with anti-GAGA factor antibodies as a loading control (Raff *et al*, 1994).

**Figure S2.**
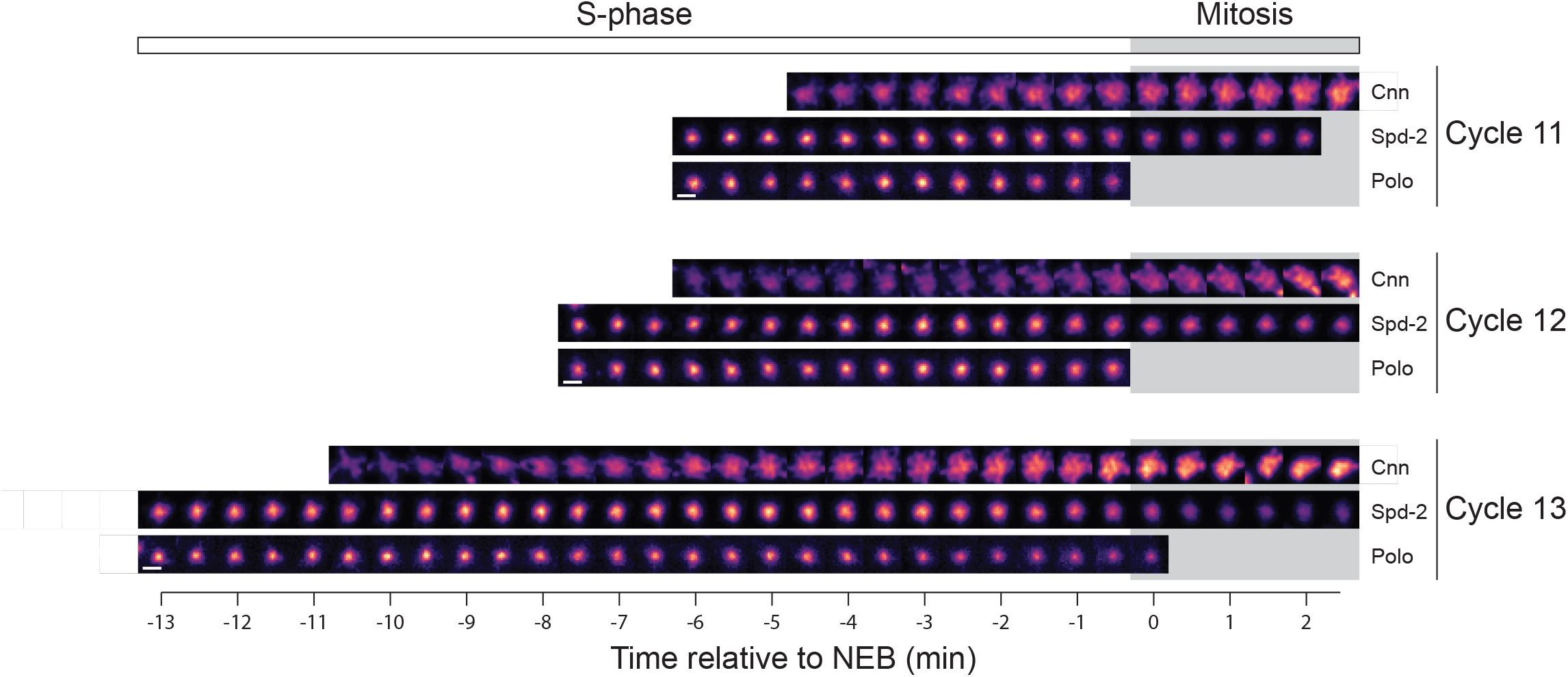
Spd-2, Polo and Cnn are recruited to assembling centrosomes with different kinetics. Images show the recruitment of either mNG-Cnn, Spd-2-GFP or Polo-GFP at an exemplar centrosome during nuclear cycles 11, 12 and 13. All images are aligned to nuclear envelope breakdown (NEB; t=0). The white parts of the graphs indicate S-phase and the grey parts mitosis. The first (most leftward) image in each series is taken when the two centrosomes associated with each nucleus at the end of mitosis have first completely separated from one another in early S-phase; because the Cnn scaffold is significantly larger than the Spd-2 or Polo scaffold, it takes longer for the two centrosomes to fully separate, so there are less images of Cnn in S-phase. Scale bar = 1μm

**Figure S3.**
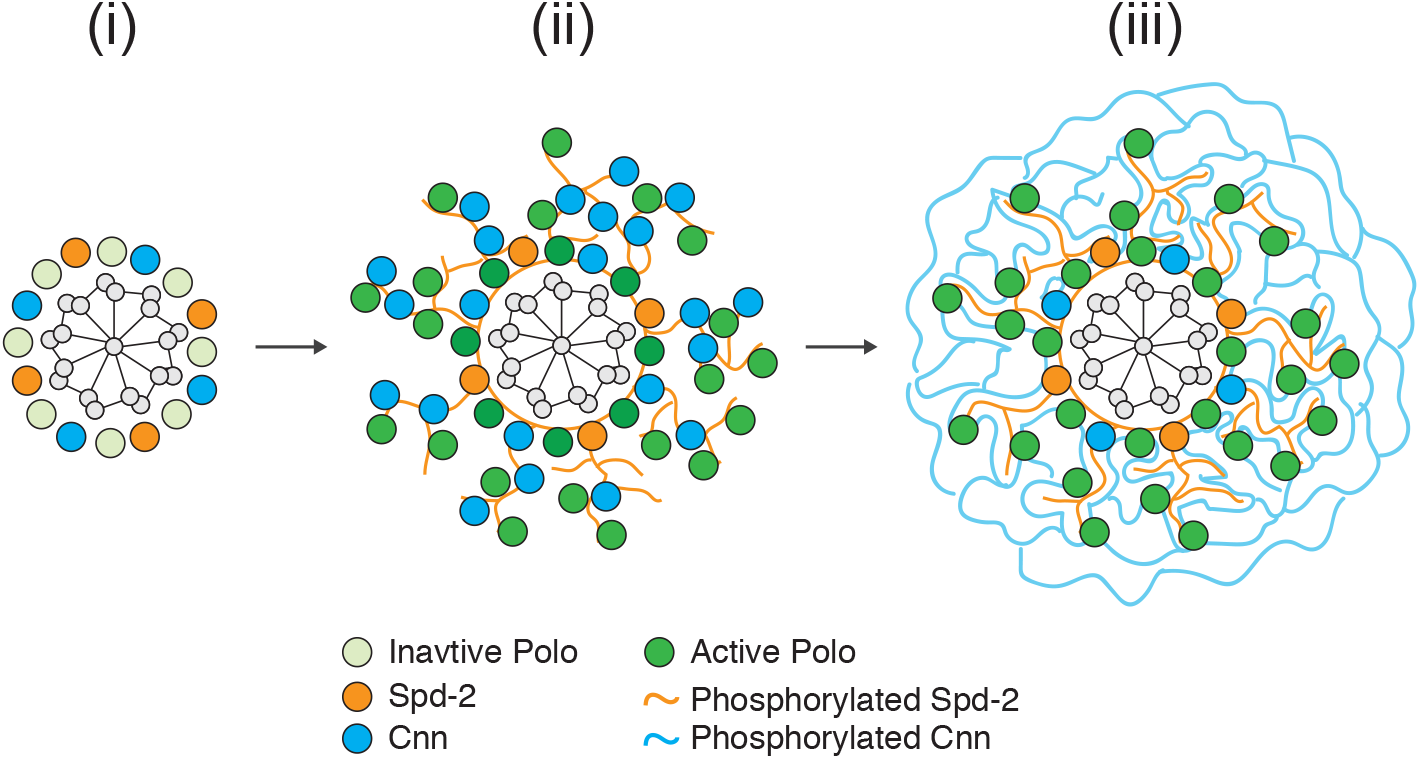
A molecular model of how Spd-2, Polo and Cnn cooperate to form a mitotic PCM scaffold. Cartoon illustrates the assembly of the Spd-2/Polo/Cnn mitotic PCM scaffold in *Drosophila*. During interphase **(i)**, Spd-2, Polo and Cnn are recruited to a toroid that surrounds the mother centriole (Fu & Glover, 2012). Polo is presumably inactive, and Spd-2 and Cnn are presumably not phosphorylated. As cells prepare to enter mitosis **(ii)**, Polo is activated at the centriole and the centrosomal Spd-2 becomes phosphorylated, allowing it to assemble into a scaffold that can flux outwards away from the centriole. The phosphorylated Spd-2 scaffold (equivalent to *S*\* in Figure 2B) is structurally weak, but it can recruit Polo—via phosphorylated S-S(P)/T(P) motifs (Alvarez-Rodrigo et al, 2019)— and also Cnn (Conduit et al, 2014b) to form the more stable 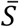 scaffold depicted in Figure 2B. The Polo recruited by Spd-2 is activated and can phosphorylate Cnn, allowing Cnn to assemble into a strong macromolecular scaffold (*C*\* in Figure 2B) (Conduit et al, 2014a; Feng et al, 2017). Cnn itself cannot recruit more Spd-2 or Polo, but it stabilises the expanding Spd-2 scaffold, so allowing Spd-2 to accumulate around the mother centriole **(iii)** (Conduit et al, 2014b).

**Figure S4.**
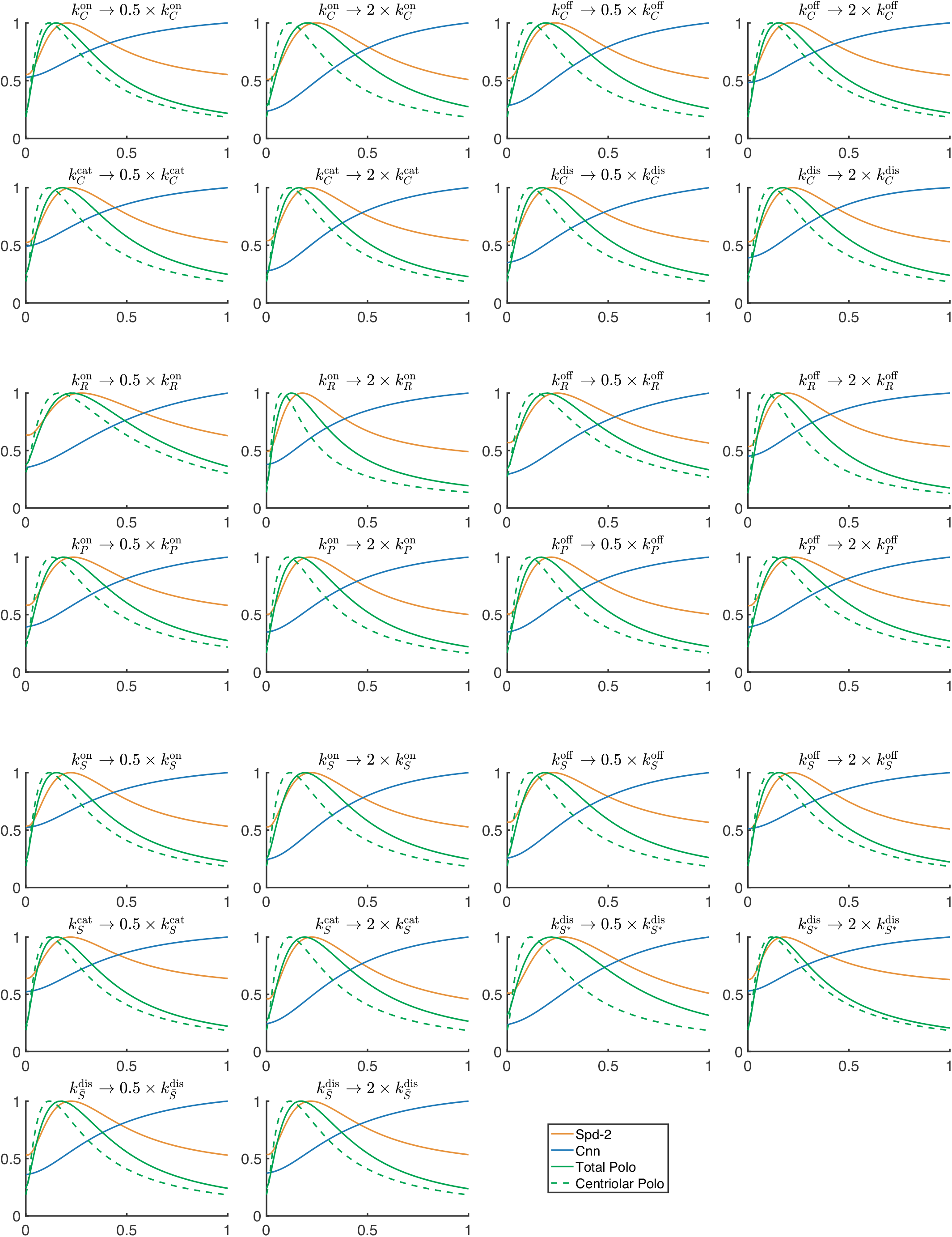
Model predictions are relatively robust to changes in parameter values. Graphs show the computed output of Model 1 and Model 2 when each of the 13 reaction rate parameters is either doubled or halved (as indicated above each graph). The qualitative behaviour of the model is consistent in all cases, demonstrating the model’s robustness in the parameter regime considered.

**Figure S5.**
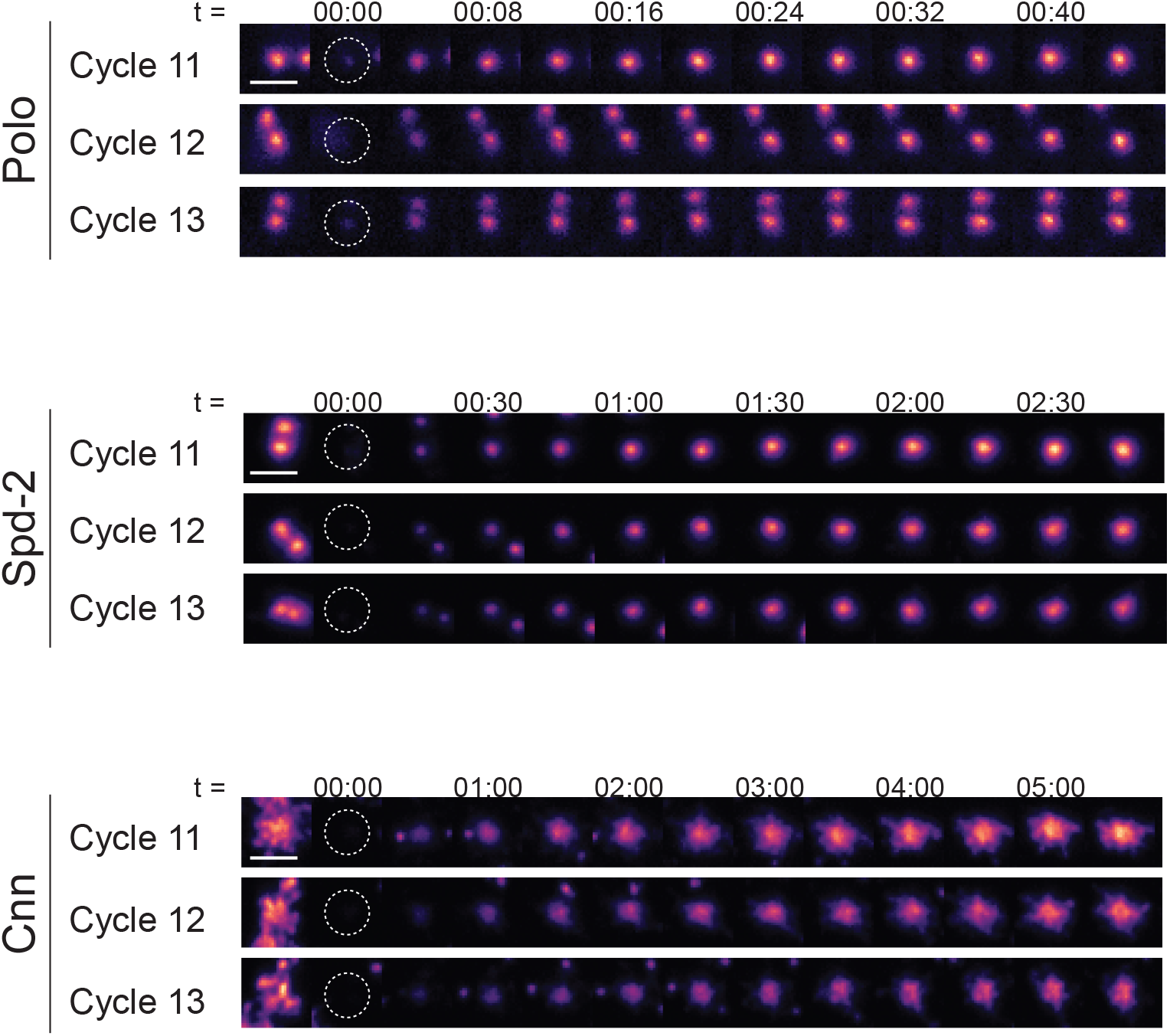
Monitoring rates of centrosomal fluorescence recovery of photobleached PCM scaffold components. Micrographs show examples of centrosomes that were fluorescently-labelled with either Polo-GFP, Spd-2-NG or NG-Cnn, photobleached at t=0, and then monitored for the subsequent recovery of fluorescence at the start of either nuclear cycle 11, 12 or 13. Time (mins:secs) is indicated above selected images (note the different time scales used for each fusion protein). Scale bar = 2μm.

## Materials and Methods

### *Drosophila melanogaster* stocks and husbandry

The *Drosophila* stocks used, generated and/or tested in this study are listed in Table 1; the precise stocks used in each experiment (and the relevant Figure) are listed in Table 2. Stocks were maintained on *Drosophila* culture medium (0.8% agar, 8% cornmeal, 1.8% yeast extract, 1% soya, 8% malt extract, 2.2% molasses, 0.14% nipagen, and 0.625% propionic acid) in 8cm x 2.5cm plastic vials or 0.25-pint plastic bottles.

**Table 1:** Drosophila stocks used in this study.

| Allele | Source |
| --- | --- |
| cnn<sup>f04547</sup> | Exelixis stock no. f04547, Exelixis Stock Centre (Harvard Medical School, Boston, MA). |
| cnn<sup>HK21</sup> | (Megraw *et al*, 1999; Vaizel-Ohayon & Schejter, 1999) |
| Ubq-NG-Cnn | Generated by Lisa Gartenmann; appears to be fully functional and rescues *cnn*<sup>-/-</sup> mutant. |
| Ubq-Spd-2-GFP | (Dix & Raff, 2007) |
| Spd-2-NG (CRISPR) | Generated for this study; appears to be fully functional and is homozygous viable and fertile. |
| Spd-2<sup>z35711</sup> | (Giansanti *et al*, 2008) |
| Spd-2<sup>G20143</sup> | (Dix & Raff, 2007) |
| Polo-TRAP-GFP | (Buszczak *et al*, 2007); appears to not be fully functional and is only viable as a heterozygote. |
| Ubq-Spd-2-mCherry | (Alvarez-Rodrigo *et al*, 2019) |
| ana1<sup>mecB</sup> | (Blachon *et al*, 2009; Avidor-Reiss *et al*, 2004) |
| Ubq-Ana1-mCherry | (Alvarez-Rodrigo *et al*, 2020) |
| Ubq-Ana1-S34T-mCherry | (Alvarez-Rodrigo *et al*, 2020) |
| Ubq-Spd-2-S16T-mCherry | (Alvarez-Rodrigo *et al*, 2019); described in this previous publication as Spd-2-CONS. |

**Table 2:**
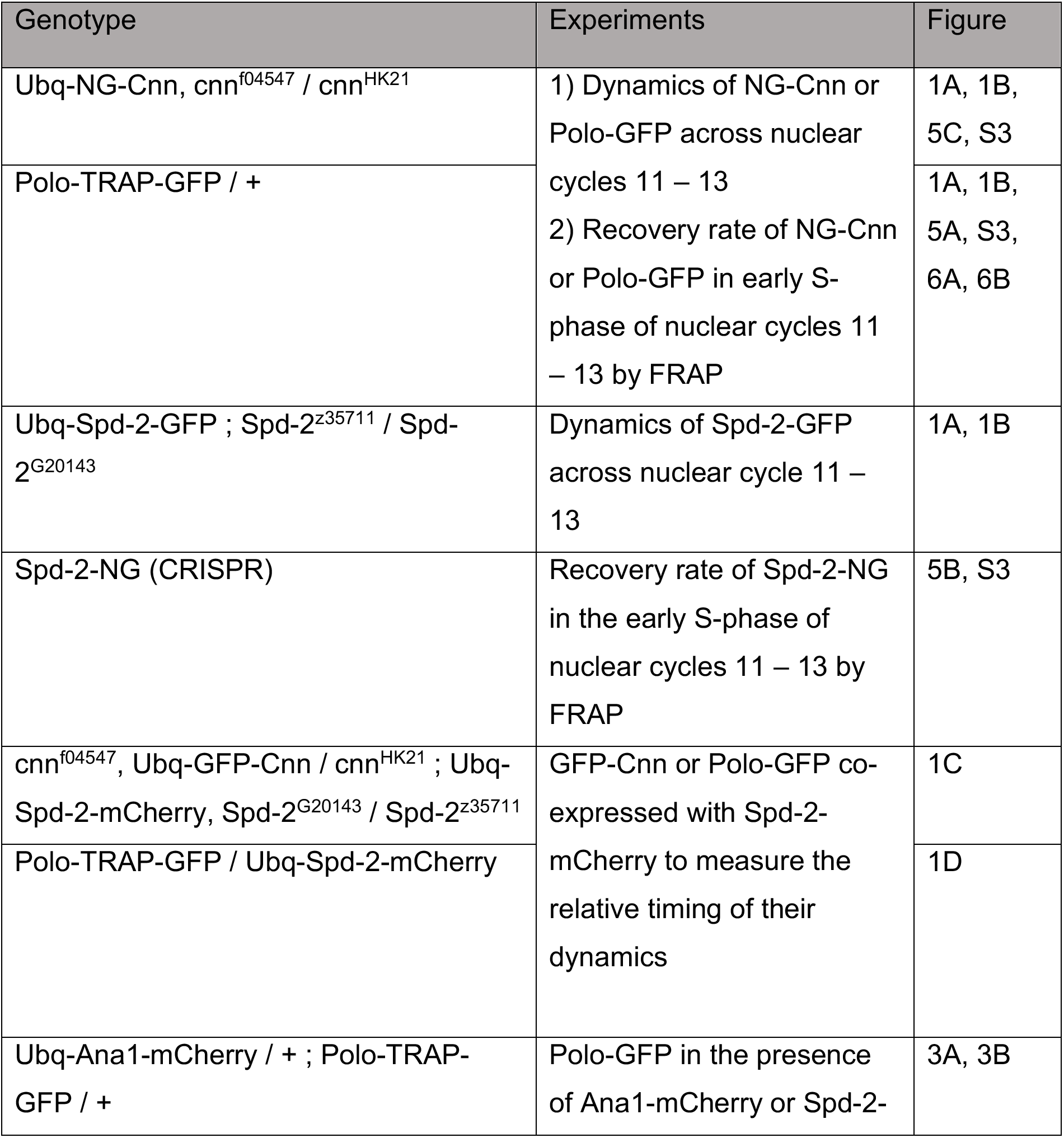
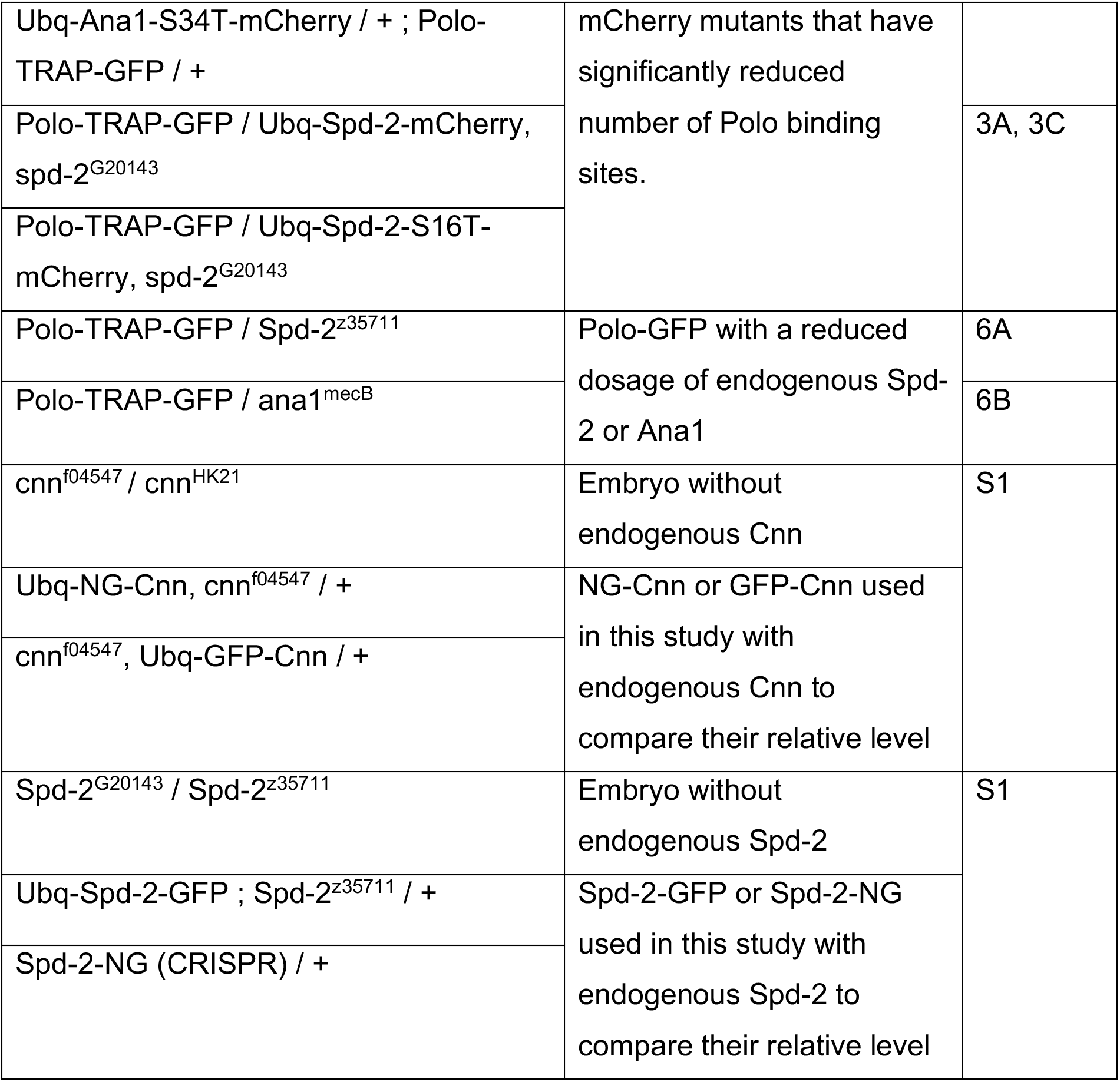
*Drosophila* stocks used in specific experiments.

### CRISPR/Cas9-mediated generation of Spd-2-mNG knock-in fly line

A single guide RNA (sgRNA) and donor plasmid for homology-directed repair (HDR) were generated respectively, injected into Cas9-expressing CFD2 embryos and screened as previously described (Port *et al*, 2014). Briefly, the sgRNA target sequence was selected that was close to the insertion site using a sgRNA design algorithm as described previously (Gratz *et al*, 2014), and cloned in pCFD3 plasmid (Port *et al*, 2014). The donor plasmid containing tandem DNA sequence of 1 kb upstream homology arm, linker plus mNeonGreen sequence and 1 kb downstream homology arm was synthesized by GENEWIZ Co. Ltd. (Suzhou, China) in pUC57. To enhance the recombination process and to linearize the plasmid in vivo, the cleavage sites (sgRNA target sequence) were introduced on either side of the 3kb sequence in the donor plasmid. In addition, the sgRNA target sequences within the homology arm of the donor plasmid were mutated (without affecting the amino acid sequence) to prevent the Cas9 from cleaving within the repair template and the knock-in construct once it had been inserted into the endogenous locus in vivo. The mixture of both constructs—Guide RNA (sgRNA) and donor plasmid—was injected into Cas9-expressing CFD2 embryos (Port *et al*, 2015) by the Department of Genetics, University of Cambridge (UK). After hatching, the single flies were crossed to a balancer line (PrDr/TM6C) and screened for the positive insertion event by PCR for 2 or 3 generations. The final generation of flies was balanced, and the gene sequence containing the 3kb insertion fragment and the region flanking the insertion was sequenced.

### Embryo collections

Embryos were collected from plates (25% apple & raspberry juice, 2.5% sucrose, and 2.25% agar) supplemented with fresh yeast suspension. For imaging experiments, embryos were collected for 1h at 25°C, and aged at 25°C for 45–60 min. Embryos were dechorionated by hand, mounted on a strip of glue on a 35-mm glass-bottom Petri dish with 14 mm micro-well (MatTek), and desiccated for 1 min at 25°C before covering with Voltalef grade H10S oil (Arkema). Embryo collections for western blotting experiments were performed as described previously (Novak *et al*, 2014).

### Immunoblotting

Immunoblotting analysis to estimate protein expression level was performed as described previously (Aydogan et al., 2018). The following primary antibodies were used: rabbit anti-Spd-2 (1:500) (Dix & Raff, 2007), rabbit anti-Cnn (1:1000) (Lucas & Raff, 2007), and rabbit anti-GAGA factor (1:500) (Raff *et al*, 1994). HRP-conjugated donkey anti-rabbit (NA934V lot:17876631, Cytiva Lifescience) secondary antibodies were used at 1:3000.

### Spinning disk confocal microscopy

Images of embryos were acquired at 23°C using a PerkinElmer ERS spinning disk confocal system mounted on a Zeiss Axiovet 200M microscope using Volocity software (PerkinElmer). A 63X, 1.4NA oil objective was used for all acquisition. The oil objective was covered with an immersion oil (ImmersolT 518 F, Carl Zeiss) with a refractive index of 1.518 to minimize spherical aberration. The detector used was a charge-coupled device (CCD) camera (Orca ER, Hamamatsu Photonics, 15-bit), with a gain of 200 V. The system was equipped with 405nm, 488nm, 561nm, and 642 solid-state lasers (Oxxius S.A.). All red/green fluorescently tagged samples were acquired using UltraVIEW ERS ‘Emission Discrimination’ setting. The emission filter of these images was set as followed: a green long-pass 520nm emission filter and a red long-pass 620nm emission filter. For dual channel imaging, the red channel was imaged before the green channel in every slice in a z-stacks. For Fluorescent Recovery after Photobleaching (FRAP) experiments, circular regions of interests (ROI) of diameter 4 μm were defined around selected centrosomes of interest (multiple centrosomes were often selected from a single individual embryo). A 488 nm laser at 50% laser power was used to FRAP each sample in 10 iterations over a period of 2secs. 0.5-μm z-sections were acquired, with the number of sections, time step, laser power, and exposure depending on the experiment.

### Data analysis

Raw time-series from imaged embryos were imported into Fiji. The photobleaching of raw time-series images was corrected using the exponential decay algorithm and images were z-projected using the maximum intensity projection function. The background was estimated and corrected by a uneven illumination background correction (Soille, 2004). The centrosomes were tracked using TrackMate (Tinevez *et al*, 2016). A custom Python script was then used to appropriately threshold and extract the fluorescence intensities of all of the tracked centrosomes as they changed over time in each individual embryo. To extract the features of the Spd-2 and Polo oscillations we measured the *initial intensity* of the centrosomes as they first separated in early S-phase and their *maximum intensity* at the oscillation peak; the time between these points represented the *growth period*, while the *growth rate* was calculated as: (*maximum intensity* – *initial intensity)/growth period*. To extract these features for Cnn, several mathematical models were fit to the data from each embryo, and the model that best fit the majority of the embryos was then applied to all embryos: *linear increase* (Cycle 11); *linear increase + plateau* (Cycle 12); *linear increase + linear decrease* (Cycle 13) (Table 3). The average *initial intensity, maximum intensity*, *growth period* and *growth rate* were then calculated from the fitted data for each embryo.

**Table 3:**
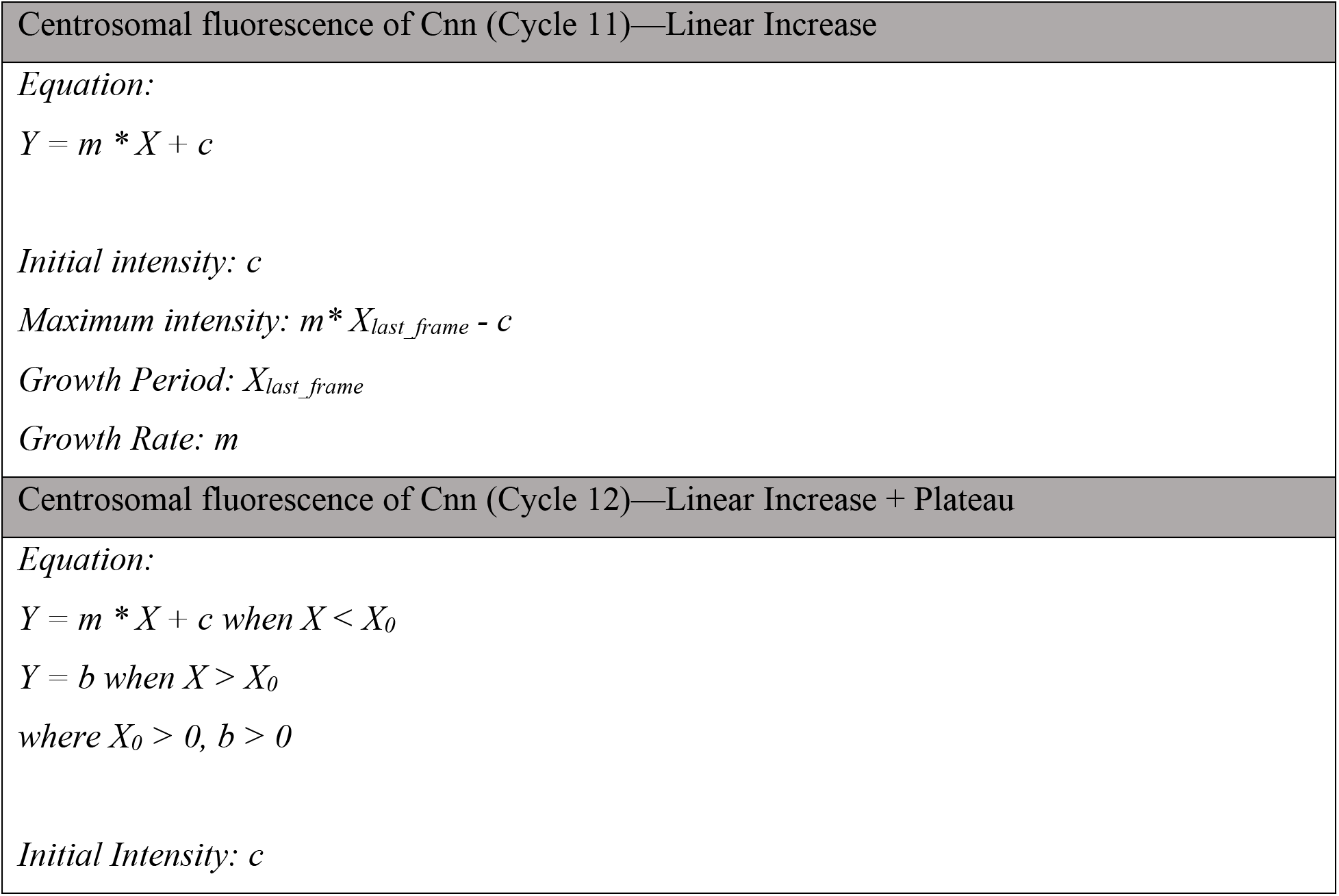
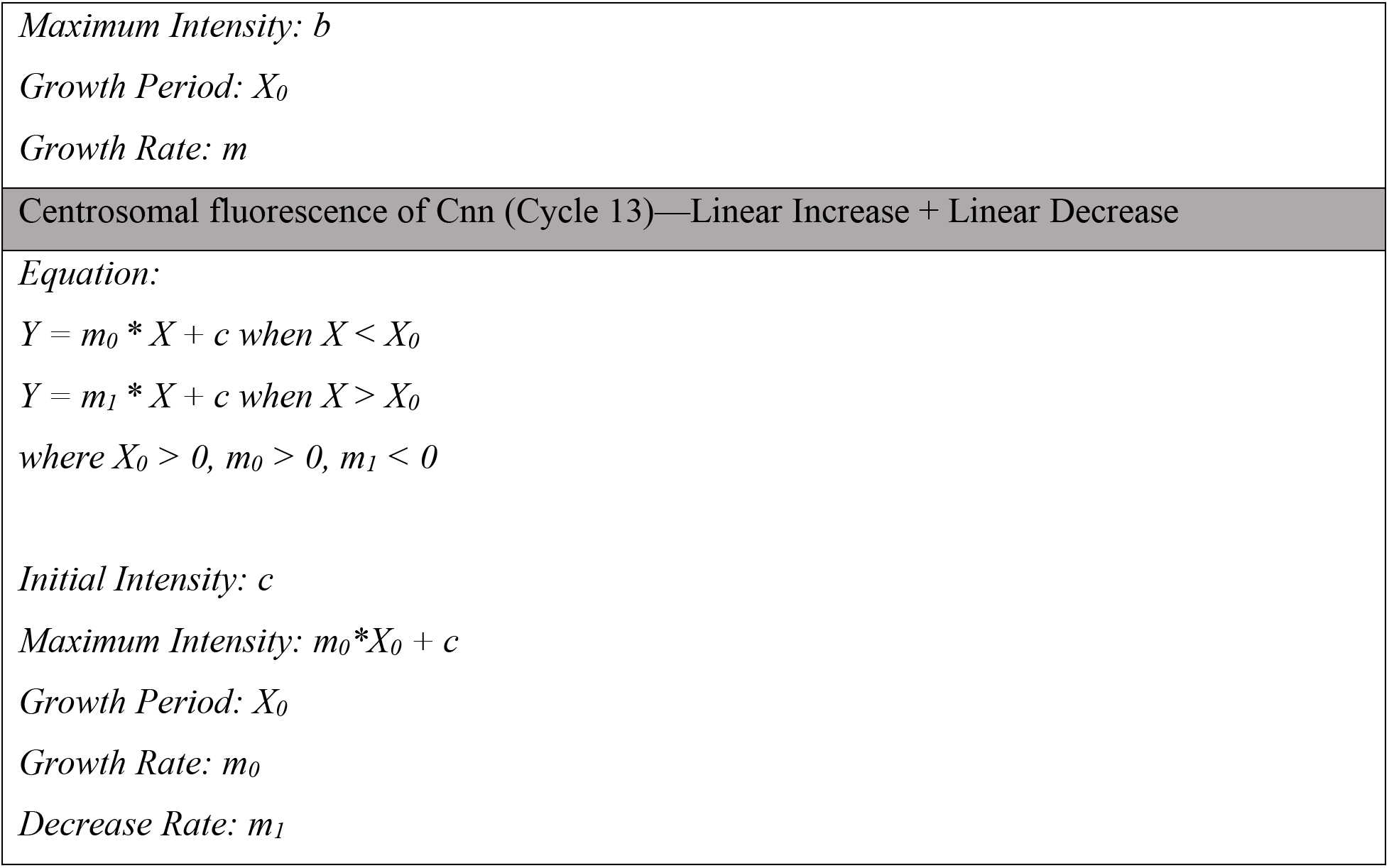
Models used for feature extraction of the Cnn data.

| Centrosomal fluorescence of Cnn (Cycle 11)—Linear Increase |
| --- |
| *Equation:*<br>$Y = m * X + c$<br>*Initial intensity:* $c$<br>*Maximum intensity:* $m * X_{last\_frame} - c$<br>*Growth Period:* $X_{last\_frame}$<br>*Growth Rate:* $m$ |
| Centrosomal fluorescence of Cnn (Cycle 12)—Linear Increase + Plateau |
| *Equation:*<br>$Y = m * X + c \text{ when } X < X_0$ $Y = b \text{ when } X > X_0$<br>*where* $X_0 > 0, b > 0$<br>*Initial Intensity:* $c$ |
| *Maximum Intensity: $b$*<br>*Growth Period: $X_0$*<br>*Growth Rate: $m$* |
| Centrosomal fluorescence of Cnn (Cycle 13)—Linear Increase + Linear Decrease |
| *Equation:*<br>$Y = m_0 * X + c$ when $X < X_0$<br>$Y = m_1 * X + c$ when $X > X_0$<br>where $X_0 > 0, m_0 > 0, m_1 < 0$<br>*Initial Intensity: $c$*<br>*Maximum Intensity: $m_0 * X_0 + c$*<br>*Growth Period: $X_0$*<br>*Growth Rate: $m_0$*<br>*Decrease Rate: $m_1$* |

For FRAP analysis, a tight bounding box was manually drawn around each centrosome (see tutorial in the Github repository of this publication), and the box was linked across multiple frames using a custom Python script. In experiments where the centrosomes organized by the old mother centriole and new mother centriole (OM and NM centrosomes, respectively) were tracked independently, two centrosomes with the shortest inter-centrosomal distance at the start of S-phase and within a preset distance threshold were annotated as a pair. The brighter centrosome in a pair was annotated as the OM while the dimmer one was annotated as NM (Conduit *et al*, 2010; Novak *et al*, 2014). The link to these custom Python scripts can be found on Github under the folder “Data analysis” (https://github.com/SiuShingWong/Wong-et-al-2021).

### Mathematical model of PCM scaffold assembly kinetics (*Model 1*)

We assume that centriolar Spd-2 receptors, *R_S_*, are able to convert cytoplasmic Spd-2, *S*, into an unstable Spd-2 scaffold, *S*\*, via the complex 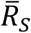. The on, off, and catalytic conversion rates of this process are 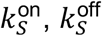, and 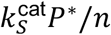, respectively, where *P*\*(*t*) is the total amount of active Polo at time *t* and *n*(*t*) is the number of centrioles in the embryo at time *t* (which will double after every cycle), so that *P*\*/*n* describes the amount of active Polo at each centriole. In the first instance, we do not attempt to model *P*\*(*t*) but instead treat it as a given function which we use as an external stimulus for the system. Although *S*\* is unstable, it can recruit cytoplasmic Cnn, *C*, and cytoplasmic Polo, *P*, to form the more stable complex 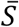, which can phosphorylate *C* to convert it into a stable Cnn scaffold form *C*\*. The on, off, and catalytic conversion rates of this process are 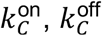 and 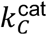, respectively. The two scaffold forms of Spd-2 have disassembly rates 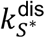 and 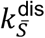, respectively, while the disassembly rate of the Cnn scaffold is given by 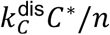, as we assume that this disassembly rate is proportional to the size of the Cnn scaffold. This assumption is based on our previous observation that at the start of S-phase the old mother (OM) centrosome organises a larger Cnn scaffold than the new mother (NM), but the two scaffolds ultimately grow to the same size by the end of S-phase (Conduit *et al*, 2010). As the Cnn incorporation rate is the same at OM and NM centrosomes (Conduit *et al*, 2010; S.S.W, *unpublished observations*) we infer that the rate of loss of Cnn during S-phase must be larger at the OM, indicating that the larger the Cnn scaffold, the larger the rate of Cnn loss.

The previous description can be summarised as a system of reactions

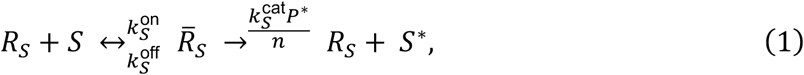

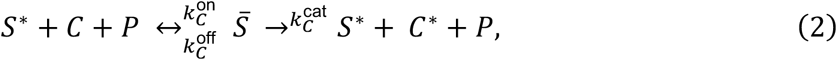

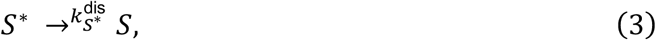

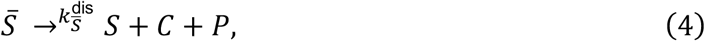

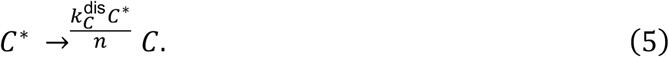

For simplicity, we assume that cytoplasmic species diffuse sufficiently fast in the embryo that we may treat these variables as spatially homogeneous, and therefore we neglect spatial effects from the model. By imposing the law of mass action, we derive the following system of four ordinary differential equations (note that the explicit dependence in the dependent variables on time has been dropped)

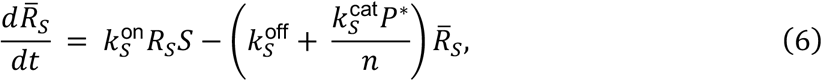

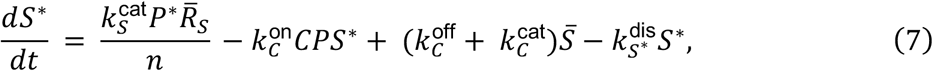

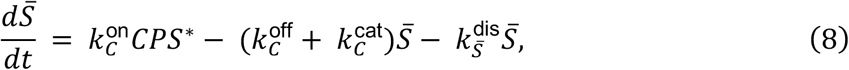

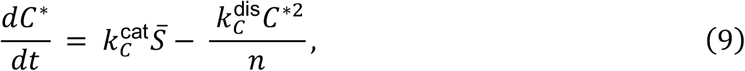

where the PCM quantities, 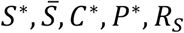 and 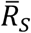 are defined as the total number of the corresponding species in the embryo (i.e. dimensionless units), and the cytoplasmic quantities, *S, C*, and *P*, are defined as the volumetric concentration of the corresponding species (i.e. units m^−3^). We assume, for simplicity, that the embryo is a closed system which implies that the total amount Spd-2 (*S*_0_) and Cnn (*C*_0_) in the embryo is conserved. Further, since the total amount of Polo in the system (*P*_0_) is large (Casas-Vila *et al*, 2017) we treat cytoplasmic Polo as a prescribed constant unaffected by absorption into the scaffold. Finally, we assume that the total number of Spd-2 receptors in the embryo is proportional to the number of centrioles. These constraints read

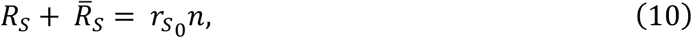

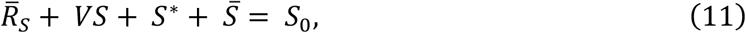

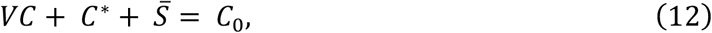

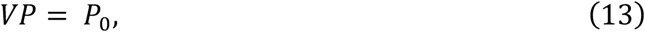

where *r*_*S*_0__ is the total number of receptors per centriole and *V* is the volume of the embryo.

These equations describe the total amount of each species in the embryo. However, it is useful to describe the model on a per-centriole basis. We do this by defining the auxiliary (lower case) per-centriole variables: 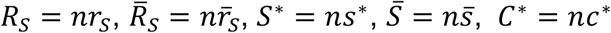, and *P*\* = *np*\*. In terms of these variables, our system reads

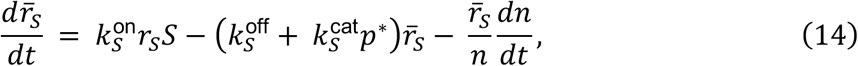

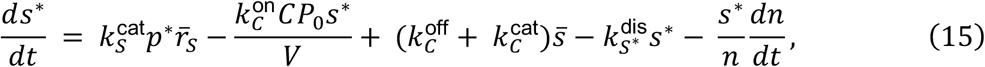

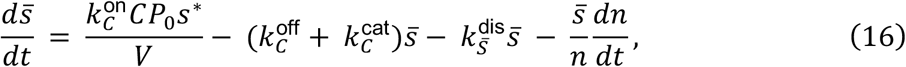

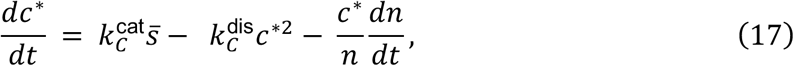

subject to

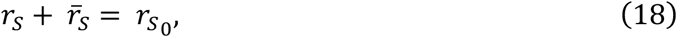

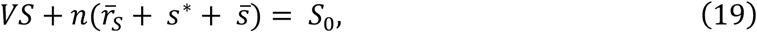

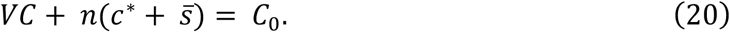

While equations (14) – (20), subject to the appropriate initial conditions, are sufficient to describe the system, it is convenient for its mathematical analysis to instead formulate the model in terms of “dimensionless” variables. Through this process, we determine the dimensionless parameter groups (e.g. the ratio of the reaction rates to the cell cycle timescale) which govern the dynamics of the system, which in turn enables us to simplify the system and reduce the number of independent variables in the model. We non-dimensionalise the system by using the following scalings

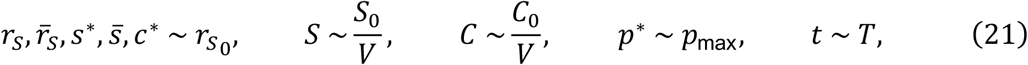

where *T* is the typical period of the cell cycle, and *p*_max_ is the maximum amplitude of the imposed Polo activity. In terms of dimensionless variables, the model reads

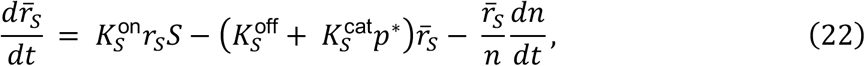

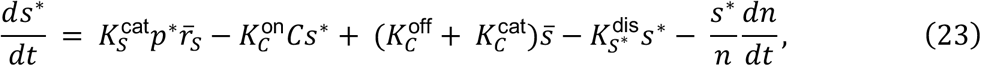

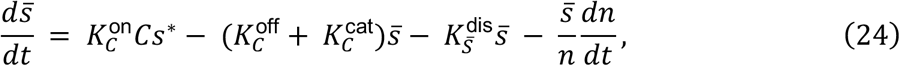

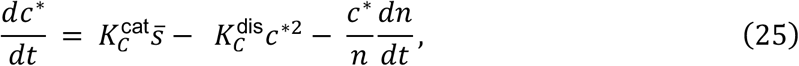

subject to

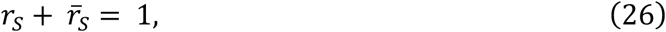

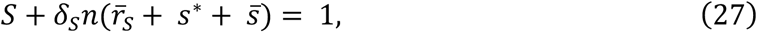

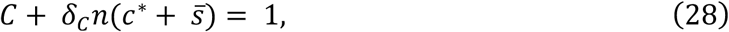

where

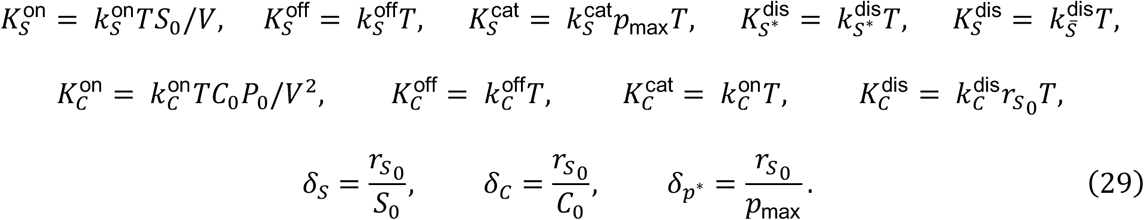

Given a solution to this system, the total size of the Spd-2 and Cnn scaffolds and total amount of active Polo surrounding each centriole are given by

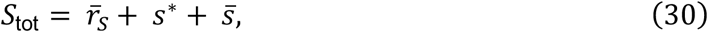

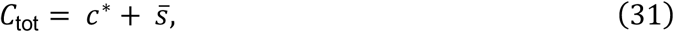

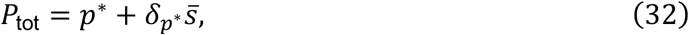

where *S*_tot_, *C*_tot_, and *P*_tot_ are dimensionally scaled with *S*_0_, *C*_0_, and *p*_max_, respectively. To allow us to compare accurately the output from our models to the experimental data we first determined reasonable initial conditions, as the centrosomes in our experiments are already initially associated with some PCM (that was acquired in the previous cycle) at the start of S-phase. To do this, we first solve (22) – (28) subject to the initial conditions 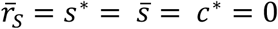. (i.e. no Spd-2 or Cnn scaffold is assembled around the centriole). Since the system is approximately cyclic, we then use the final values output by this solution, 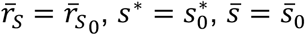, as our new initial values. Since the Cnn scaffold divides and partially breaks away during centriole separation, we cannot impose the cyclic condition on Cnn. However, since the output of *c*\* is *O*(1), this suggests that an *O*(1) input is consistent with our model and therefore we set *c*\* = 1 as our initial condition. In this way, the centrioles in our model start the cycle already associated with some Spd-2 and Cnn scaffold that they acquired in the previous cycle, as is the case with our experimental data.

In Figure 2B, we plot the incorporation of Spd-2 and Cnn into the PCM, *S*_tot_ and *C*_tot_, over the duration of a single cycle by solving (22) – (28) subject to the initial conditions 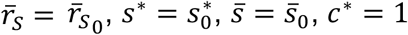, the parameter values given in **Table 4**, and the constraint that the number of centrioles is constant during the cycle, *n*≡ 1 without loss of generality. We also plot the prescribed Polo activity (i.e. the oscillation in *p*\*(*t*) that we impose on the system), 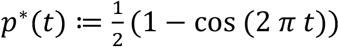, and the total Polo at the centriole, *P*_tot_. The amplitudes in all the solutions have been normalised to 1.

**Table 4:** Initial Conditions and Parameters used in Model 1.

|  |  |  |  |  |  |  |  |  |  |  |  |
| --- | --- | --- | --- | --- | --- | --- | --- | --- | --- | --- | --- |
| $K_S^{\text{on}}$ | $K_S^{\text{off}}$ | $K_S^{\text{cat}}$ | $K_{S^*}^{\text{dis}}$ | $K_{\bar{S}}^{\text{dis}}$ | $K_C^{\text{on}}$ | $K_C^{\text{off}}$ | $K_C^{\text{cat}}$ | $K_C^{\text{dis}}$ | $\delta_S$ | $\delta_C$ | $\delta_{p^*}$ |
| 20 | 200 | 100 | 20 | 1 | 100 | 100 | 50 | 0.05 | 0.001 | 0.001 | 10 |

The initial conditions and parameters used in this model are listed in Table 1. Our justification for choosing these parameter values is presented in a later Section.

We note that, in our model, the dissociation rates of the *C*\* and 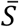 scaffolds have different functional forms. To investigate if this contributes to the different behaviour of the *C*\* and 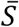 scaffolds we compare in the graphs below the model output in the case in which the exponent of the *C*\* disassembly term (□) is varied, and the case in which the disassembly rate itself is varied. This shows that *C*\* behaviour is primarily determined by the order of magnitude of the disassembly rate, rather than its exponent.

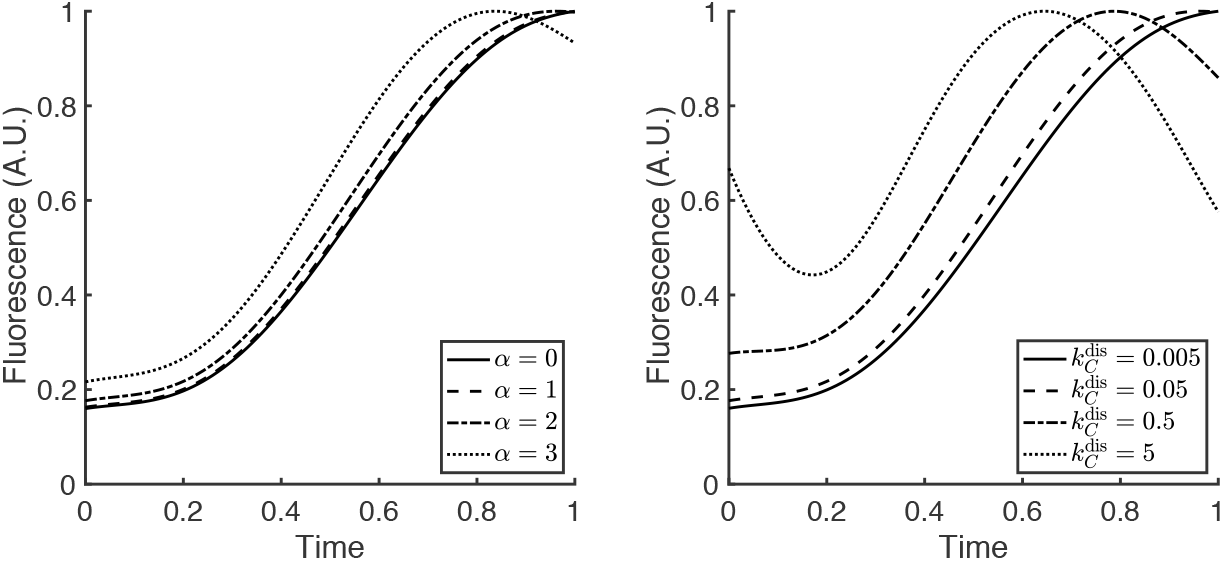

### Mathematical model of centriolar Polo activity (*Model 2*)

Model 1 assumed a given oscillation in Polo. Next, we describe a model for how such an oscillation in Polo activity might be generated by the centriole through the interaction between Polo and its receptors at the centriole surface, such as Ana1 (Alvarez-Rodrigo et al., 2021). We assume that these receptors, 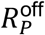, are initially inactive and unable to bind Polo. To initiate mitotic PCM assembly, the receptors are activated at a rate 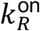 due to their phosphorylation by a protein kinase, which is most likely a Cdk/Cyclin, or a kinase that is regulated by the Cdk/Cyclins (such as Polo or Aurora A). This new form, which we denote *R_P_*, is able to bind Polo with on and off rates 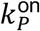 and 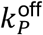, respectively, to form the complex 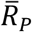. We assume that the Polo in this complex is active and able to initiate mitotic PCM assembly as described by Model 1. We also assume that this active form of Polo instigates the deactivation of the receptors at a rate 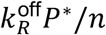. This final form, which we denote 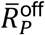, is unable to bind or activate Polo. This system likely resets itself between cycles when 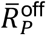 is dephosphorylated to regenerate 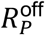, but we do not model this reset here. Finally, we assume that the reactions occurring in the PCM are the same as before, with the active centriolar Polo (in this instance given by 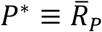) now forming part of the solution to our model. The reactions describing the generation of Polo read

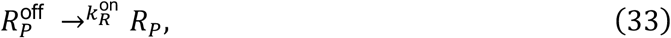

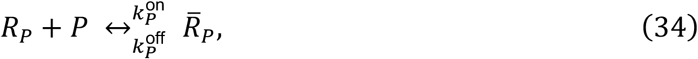

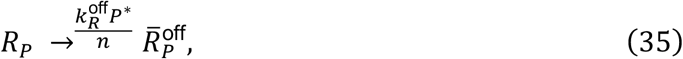

By imposing the law of mass action, we obtain the following system of ODEs,

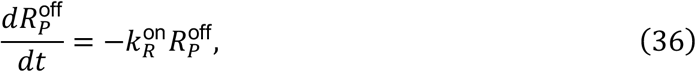

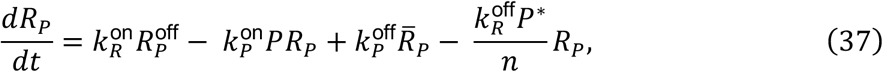

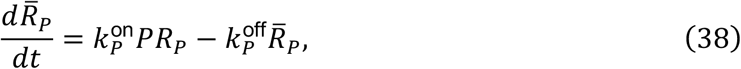

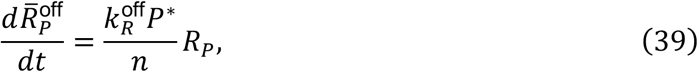

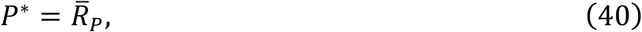

We also assume that the total number of Polo receptors at each centriole, *r*_*P*_0__, is conserved, which reads

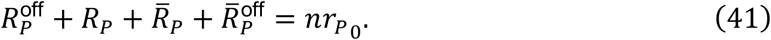

As before, we write the system in per-centriole variables, and non-dimensionalise by setting 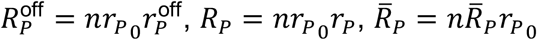, and 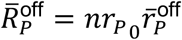, and *P*\* = *nr*_*P*_0__*p*\* so that the dimensionless model reads

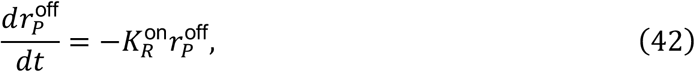

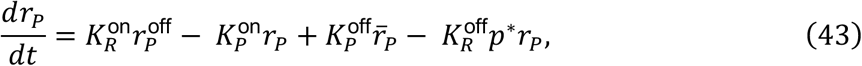

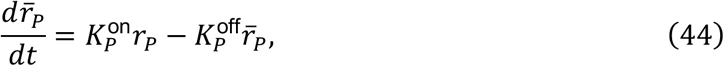

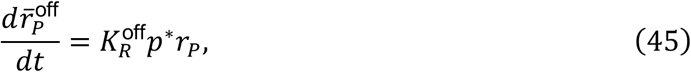

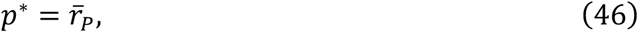

subject to

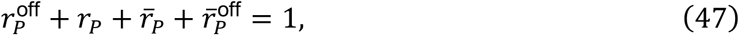

where

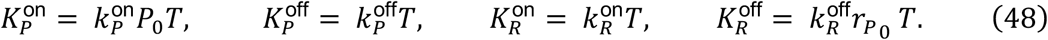

Note that we have scaled *P*\* with *r*_*P*_0__ in this instance rather than *p*_max_ since the maximum amplitude of the Polo activity is not known *a priori*., and, since the receptors generate the active Polo in this model, this is the correct scaling for *P*\*.

In this model, the total amount of Polo in the centrosome is given by

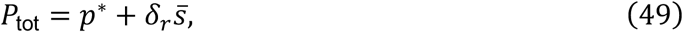

where 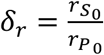.

As before, to determine the appropriate initial conditions, we first solve the model subject to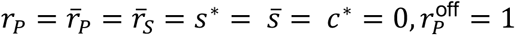 to compute the output 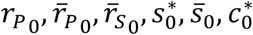. Our new initial conditions are then given by setting 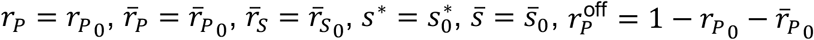.

In Figure 4A, we plot the centriolar Polo, *p*\*, the total Polo, *P*_tot_, the Spd-2 scaffold size, *S*_tot_, and the Cnn scaffold size *C*_tot_, found by solving (22) – (28) and (43) – (48) with the parameter values given in **Tables 4 and 5**. As before, to determine the appropriate initial conditions, we first solve the model subject to 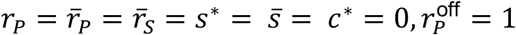 to compute the output 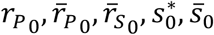. Our new initial conditions are then given by setting 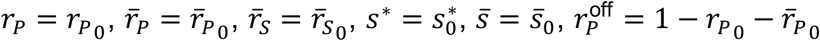, and *c*\* = 1. All solutions have been normalised.

**Table 5:** Initial Conditions and Parameters used in Model 2.

| $K_P^{\text{on}}$ | $K_P^{\text{off}}$ | $K_R^{\text{on}}$ | $K_R^{\text{off}}$ | $\delta_r$ |
| --- | --- | --- | --- | --- |
| 100 | 50 | 10 | 20 | 5 |

In Figure 6, we plot the total Polo under normal conditions (parameter values given in Tables 1 and 2) as well as half dose Ana1 (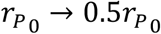, i.e. 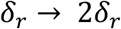 and 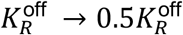) and half dose Spd-2 (*S*_0_ → 0.5*S*_0_, i.e. *δ_S_* → 2*δ_S_* and 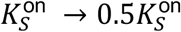). All solutions have been normalised with respect to the wild type solution.

The MATLAB scripts to recapitulate the findings of all models can be found on Github under the folder “Mathematical modelling” (https://github.com/SiuShingWong/Wong-et-al-2021).

### Justification of parameter values

We drew on a number of sources to estimate the relative magnitudes of the dimensionless reaction rate parameters (Table 4 and 5) for the models depicted schematically in Figures 2B and 4A.

Due to the rapid fluorescence recovery rates observed in FRAP experiments (Conduit et al., 2010, 2014; Feng et al., 2017) (Figure 5), it follows that the Polo reaction rates, 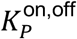 Spd-2 reaction rates, 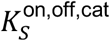, and Spd-2 scaffold disassembly rate, 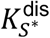, are large relative to the cell cycle timescale. Furthermore, due to the large size and rapid construction rate of the Cnn scaffold, it follows that 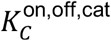, are also large. By contrast, since FRAP data shows that the fluorescence level of the Cnn scaffold fails to fully recover even over an entire nuclear cycle (Figure 5), it follows that the Cnn scaffold disassembly rate, 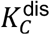, is small by comparison with the cell cycle timescale. In order to quantify the modelling assumption that the 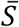 scaffold is more stable than the *S*\* scaffold, we prescribe that 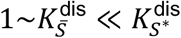.

We expect that the catalytic conversion, i.e. activation and subsequent release, of Spd-2 and Cnn is a more complex process than unbinding alone, so we assume the catalytic conversion rates are slower than the off rates. However, for simplicity, we assume that these rates are all of a similar order of magnitude. To ensure that the Spd-2 scaffold can convert Cnn into a scaffold before it disassembles, we also assume that the Spd-2 disassembly rate is less than the catalytic conversion rate of Cnn.

We make the additional assumption that 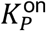 is larger than 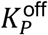 since the amount of Polo in the embryo is large (Casas-Vila *et al*, 2017) and 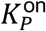 is proportional to *P*_0_. By contrast, we assume that 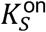 is smaller than 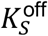 and 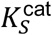 as the amount of Spd-2 in the embryo is comparatively small (Casas-Vila *et al*, 2017) and 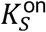 is proportional to *S*_0_. Our own Fluorescence Correlation Spectroscopy (FCS) data indicate that Polo is present in the cytoplasm at 3-5X higher levels than Spd-2 or Cnn (Thomas Steinacker, *personal communication*).

Since the Polo receptors are required to both activate and deactivate in a single cycle, it follows that 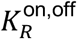 are sufficiently large (i.e. greater than order unity) that the receptors have time to reset, but not so large that the resetting is instantaneous. We therefore suppose that they are ≈ 10 for simplicity.

These assumptions may be combined to read

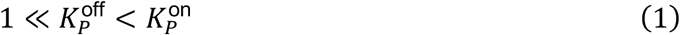

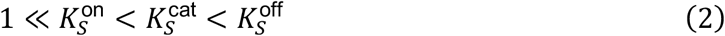

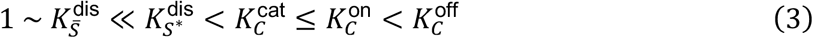

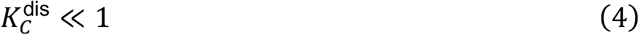

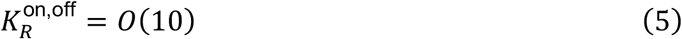

Since the total amount of Spd-2 and Cnn in the embryo likely greatly exceeds the number of Spd-2 receptors at the centriole, we prescribe *δ_S_* ≪ 1. Furthermore, we observe through Fluorescence Correlation Spectroscopy (FCS) analysis that the total amount of Spd-2 and Cnn in the embryo are similar (Thomas Steinacker, *personal communication*) and therefore *δ_S_* ≈ *δ_C_*. On the other hand, since the total amount of Polo bound to the centriole cannot exceed the number of receptors, it follows that *δ*_*P*\*_ > 1. Finally, since we are unable to determine the relative sizes of *r*_*P*_0__ and *r*_*S*_0__, we suppose that *δ_r_* = *O*(1) for simplicity. Hence, these parameters satisfy

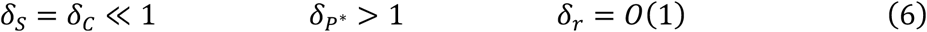

The assumptions outlined above lay the foundation for the parameter values we have chosen. However, it worth noting that the characteristic behaviour of the response curves is unaltered by doubling or halving any of these values, and therefore the particular regime we analyse in this manuscript is robust to variation in the parameters.

### Statistical analysis

The details of statistical tests, sample size, and definition of the centre and dispersion are provided in individual Figure legends.

